# CD38^+^ Endothelial Remodeling Defines Spatially Diverse Vasculopathy Programs in Rapidly Advancing Oral Inflammation

**DOI:** 10.1101/2025.07.29.667333

**Authors:** Quinn T. Easter, Khoa L. A. Huynh, Camila Schmidt Stolf, Jialiu Xie, Bruno F. Matuck, Akira Hasuike, Zabdiel Alvarado-Martinez, Zhaoxu Chen, Apoena Aguiar Ribeiro, Nivedita Pareek, Andrea M. Azcarate-Peril, Di Wu, Renato Casarin, Kang I. Ko, Jinze Liu, Kevin M. Byrd

## Abstract

Oral inflammatory diseases affect nearly half of the global population. Among them, newly defined peri-implantitis and high-grade periodontitis represent rapidly advancing inflammatory disease types, marked by relatively rapid tissue destruction. Despite their prevalence, the cell mechanisms and spatial architecture driving this severity remain poorly understood. Focusing first on peri-implantitis versus low- and moderate-grade periodontitis, we applied microbial profiling, single-cell RNA sequencing (scRNA-seq), and spatial proteomics (sp-proteomics) to uncover shared pathogenic programs linked to accelerated niche breakdown. Furthermore, to preserve spatial fidelity, each tissue was anatomically orientated along the tooth– or implant– epithelial interface, analogous sites of disease origination. Laser capture microdissection followed by microbiome analysis of unique tissue compartments revealed reduced bacterial load and diversity in peri-implantitis stroma. We then expanded our version-1 Human Periodontal Atlas by integrating newly generated peri-implantitis scRNAseq data (36-total samples; 121395-cells), revealing widespread transcriptional alterations, including oxidative stress, hypoxic, and NAD^+^ metabolism-associated signatures, primarily in a subpopulation of *TNFRSF6B^+^*/*ICAM1^+^*post-capillary venules. We then performed high-resolution sp-proteomics (15-total samples; 337260-cells) and analyzed VEC states and associated neighborhoods via *AstroSuite* using newly developed tri-wise spatial analysis. This revealed CD34^+^-VEC loss and CD38^+^-VEC expansion almost exclusively in peri-implantitis. We extended this analysis to high-grade periodontitis. Mucosal biopsies from four lesion-affected and four unaffected sites within the same individuals (1:1 matched; 8-samples; 225137-cells) again demonstrated spatially restricted CD38^+^-VEC remodeling exclusively in affected tissues, with similar vasculopathy front patterning. The findings nominate spatially distinct vasculopathy patterning as a hallmark of rapidly advancing oral inflammation and a targetable therapeutic axis.

## INTRODUCTION

Mucosal inflammatory diseases of the oral cavity affect billions of individuals globally and remain a major obstacle to precision oral medicine^1,2^. Their clinical heterogeneity and limited molecular characterization have long hindered the development of targeted interventions^3–5^. While the roles of immune cells in chronic inflammation are well-established^6^, the contributions of structural cell types to these inflammatory diseases i.e., epithelial and stromal cells, remain less clearly defined. These cells form the foundation of the tissue microenvironment by establishing physical barriers and regulating nutrient exchange; however, emerging spatial multiomic approaches have illuminated how structural cells also coordinate immune cell recruitment and local tissue regulation^7–9^. This growing body of work suggests that structural cell dysfunction may not simply result from inflammation but may actively initiate and sustain pathogenic immune programs, including structural cell inflammatory memory programs^10–13^. However, despite these insights, the spatial and transcriptional reprogramming of structural cells during chronic disease progression, particularly in rapidly advancing lesions, remains poorly understood.

Among mucosal inflammatory diseases, those that follow a rapidly destructive course offer key insights into the mechanisms that drive unchecked tissue breakdown. Rapidly advancing oral inflammatory diseases, such as recently-defined peri-implantitis^14^ and high-grade periodontitis (Stage III/Grade C; aka “generalized rapidly advancing periodontitis”)^15,16^, provide a valuable opportunity to investigate how structural and immune cell dysfunction contribute to this pathology. Despite sharing clinical hallmarks with more indolent forms of periodontitis, these two lesions exhibit accelerated destruction of both soft and hard tissues, often in shorter timeframes and with limited therapeutic responsiveness^17,18^. These conditions disproportionately compromise structural integrity and lead to significant clinical morbidity, yet the molecular distinctions that separate them from slower-advancing forms remain poorly resolved.

Importantly, they arise in anatomically adjacent but biologically distinct niches, implant versus tooth, offering a unique comparative framework for exploring how local microenvironments influence inflammatory programming and tissue breakdown. As such, these lesions serve as ideal models to study how spatially compartmentalized structural cells contribute to chronic inflammation and rapid mucosal degradation.

To interrogate the structural and spatial programs underlying rapid mucosal tissue destruction, we developed a multi-cohort, cross-disease framework integrating histopathology, laser capture microbiome profiling, single-cell RNA sequencing (scRNA-seq), and high-resolution spatial proteomics. All tissue samples were anatomically oriented along the tooth or implant interface to preserve spatial fidelity and enable direct comparisons of disease initiation sites. We first focused on peri-implantitis versus periodontitis, demonstrating that peri-implantitis lesions in this cohort are larger and display altered stromal matrix deposition. We then developed a novel laser capture microdissection followed by whole-genome microbial sequencing protocol, revealing that, despite exhibiting more severe clinical and histological destruction, peri-implantitis lesions paradoxically contained lower microbial quantity and diversity than either periodontitis or healthy tissues, especially in the stroma, highlighting the importance of host-driven mechanisms in lesion expansion.

To further uncover disease-specific cell states and transcriptional programs in a rapidly progressive inflammatory disase, we expanded our v1 Human Periodontal Atlas by integrating newly generated scRNA-seq data from peri-implantitis samples, totaling 121,395 cells^8,19^. This integrated analysis revealed widespread transcriptional reprogramming across nearly all major cell types, including epithelial cells, fibroblasts, and vascular endothelial cells (VECs). Notably, VECs were markedly increased in number within the disease-associated stroma, forming a distinct vascular compartment. This population exhibited upregulation of signaling pathways related to EGFR, insulin, leptin, IL-6, and B cell receptor activation. The most striking alterations were found in a subpopulation of *TNFRSF6B*^+^/*ICAM1*^+^ post-capillary venules, which showed strong enrichment for oxidative stress, hypoxia, endothelial-to-mesenchymal transition (EndoMT), and NAD^+^-related metabolic stress pathways, implicating this subset in the inflammatory vasculopathy unique to peri-implantitis.

To validate and spatially contextualize these findings, we applied high-resolution spatial proteomics (15-total samples; 337260-cells) and performed integrated analysis of VECs, immune cells, and their tissue cellular neighborhoods using *AstroSuite*, a containerized pipeline for multimodal spatial biology developed by our team^7,20^. For this multi-disease comparison, we developed a novel tri-wise spatial analysis framework capable of simultaneously comparing spatial distributions of cell types, cell states, and multicellular neighborhoods. Neighborhoods were computationally defined using *TACIT* and *Constellation*, two *AstroSuite* modules that infer cell types, states, and neighborhoods. This approach revealed disease-specific CD38^+^-VEC expansion and CD34^+^-VEC depletion almost exclusively in peri-implantitis. CD38^+^-VECs formed immune-vascular interaction hubs characterized by close spatial proximity to cytotoxic T cells, helper T cells, and macrophages.

Given CD38’s established role as an NAD^+^-consuming ectoenzyme that regulates intracellular NAD^+^ levels and sirtuin signaling, its upregulation in VECs suggests a metabolic shift toward NAD^+^ depletion, oxidative stress and other vasculopathy phenotypes, congruent with transcriptomic enrichment for NAD^+^ metabolic dysregulation observed in peri-implant venules. To generalize these findings, we recruited a cohort of Stage III/Grade C periodontitis subjects, profiling 1:1 matched mucosal biopsies from lesion-affected and unaffected sites within the same individuals (8-total samples; 225137-cells). Again, CD38^+^ VEC remodeling was spatially restricted to affected tissues and displayed similar immune-vascular front patterning, suggesting convergent vasculopathy programs across both rapidly progressive diseases. Together, these findings identify spatially localized endothelial remodeling, marked by CD38-mediated NAD^+^ metabolic stress, as a defining feature of rapidly advancing oral inflammation and nominate it as a potential targetable axis for intervention.

## RESULTS

### A Multimodal Strategy to Understand Accelerated Oral Inflammatory Diseases

Understanding why some oral inflammatory lesions progress rapidly while others remain indolent requires a deeper investigation into the molecular and spatial interplay between tissue-resident structural cells, immune infiltration, and microbial communities. While conventional diagnostics rely primarily on clinical and radiographic findings, these approaches fail to capture the complex biological heterogeneity underlying lesion behavior. To address this, we designed a multi-omics strategy to dissect the cellular and molecular architecture of these lesions across multiple layers of biological organization (Figure 1a). We employed histomorphometrics as well as laser capture microdissection (LCM) over epithelial and stromal compartments, followed by whole-genome microbial sequencing to assess microbial quantity and spatial distribution within healthy and inflamed tissues. We also used single-cell RNA sequencing (scRNA-seq) to define transcriptional programs of structural and immune cell types at high resolution by integrating into the human periodontal cell atlas. Finally, we leveraged high-dimensional spatial (sp)-proteomics to validate and contextualize these findings within intact tissue architecture. By integrating these orthogonal data types, we aimed to identify disease-specific cellular reprogramming, microbial-structural cell interactions, and proteomic remodeling events that together define the pathogenesis of rapidly destructive oral inflammation.

**Figure 1.**
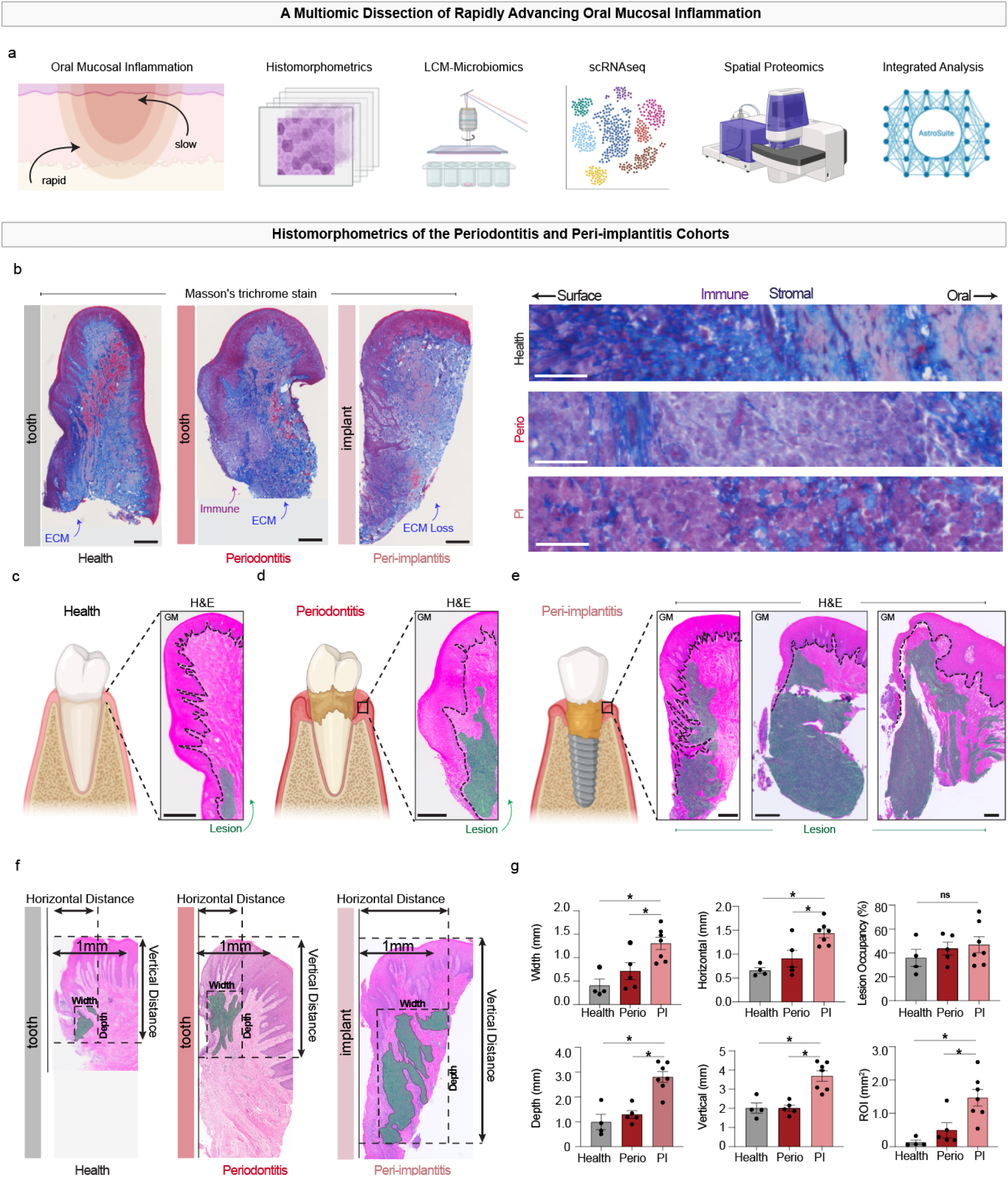
Histomorphometric analysis reveals characteristic features of periodontitis and peri-implantitis immune lesions. a) Overview of multiomic dissection of accelerated oral mucosal inflammation in this study. b) Masson’s trichrome staining of biopsies enables identification of structural-immune features, including ECM loss in peri-implantitis. H&E images of c) health, d) periodontitis, and e) peri-implantitis biopsies indicated larger immune lesions in peri-implantitis. f) Schematic of parameters used to quantify immune lesion analysis. g) Statistical analysis of length and area scales of width, horizontal and vertical length, depth, and ROI showed peri-implantitis differed from periodontitis across all metrics; lesion occupancy showed no difference. Scale bars: a), b), c), f) 300 µm; f) (inset) 20 µm; g) 50 µm. Statistical analysis: g) *p < 0.05. N = 4 (health), 5 (periodontitis), 7 (peri-implantitis). Images in a) were generated using BioRender.

### Histomorphometric Features Distinguish Peri-implantitis to Motivate Multiomics Profiling

To make head-to-head comparisons on rapidly advancing versus more slowly advancing, chronic lesions, we decided to first incorporate peri-implantitis into our growing oral and craniofacial atlasing efforts^7,21^. Peri-implantitis is a relatively recently defined inflammatory disease that affects the soft and hard tissues surrounding dental implants^22^. While implant dentistry has seen remarkable success and long-term survival rates in most patients^23^, a subset of implants develops peri-implantitis, which is characterized by mucosal inflammation, progressive bone loss, and eventual implant failure^14^. The disease presents a unique clinical challenge due to its frequently more rapid rate of progression, larger lesion size, and significantly higher densities of innate and adaptive immune cell types compared to periodontitis^24^. Despite its prevalence^25^, the biological mechanisms that drive peri-implantitis remain incompletely understood^26^.

To build on these findings and systematically interrogate the distinct tissue architectures of oral inflammatory disease, we curated a comparative cohort of human gingival biopsies from healthy individuals, patients with moderate (i.e., Stage II/III; Grade A/B) periodontitis, and patients with peri-implantitis, rigorously classified based on contemporary diagnostic criteria (Supplementary Table 1). This builds off our version 1.0 (v1) Human Periodontal Atlas, that includes health, gingivitis, and periodontitis samples, profiled using scRNAseq and/or spatial (sp)- transcriptomics, sp-proteomics, or sp-multiomic datasets^8,19^. We sought to confirm what has been known regarding peri-implantitis in our cohort to justify further investigation using an integrated multiomics approach.

We began by performing classical histological and histomorphometric analyses to assess lesion dimensions, epithelial organization, and immune-stromal interactions across conditions. Masson’s trichrome staining revealed progressive tissue disorganization from health to disease (Figure 1b; Supplementary Figure 1a). Healthy gingiva exhibited an intact extracellular matrix (ECM) and epithelial barrier; periodontitis lesions showed ECM breakdown and immune infiltration at the tooth interface. Peri-implantitis lesions, however, demonstrated the most pronounced ECM degradation, with broad, deep immune infiltration and significant stromal remodeling. High-magnification images highlighted increased intermixing of immune (purple) and stromal (blue) compartments in peri-implantitis, suggesting more extensive immune-stromal aggregation.

Hematoxylin and eosin (H&E) staining supported these observations (Supplementary Figure 1b), revealing limited immune infiltration in health (Figure 1c), largely confined to the deeper peri-crevicular and junctional zones. In contrast, both periodontitis and peri-implantitis samples exhibited substantial immune cell infiltration extending deep into the connective tissue, with lesions forming distinct localization patterns surrounding either the tooth (Figure 1d) or implant (three examples; Figure 1e). Quantitative morphometric analysis confirmed significant increases in lesion width and depth in peri-implantitis, exceeding those observed in periodontitis (Figure 1f, g). Although both disease states showed elevated lesion occupancy compared to health, only peri-implantitis demonstrated statistically significant increases across both horizontal and vertical dimensions, reinforcing its designation as a rapidly destructive lesion. Notably, even at this histological level, scattered vascular-like structures were more frequently observed throughout the peri-implantitis lesion, particularly within inflamed stromal regions (Figure 1b). These early histological cues confirmed known observations yet pointed toward increased vascular involvement, which justified deeper interrogation of the stromal compartment as a potentially overlooked contributor to lesion expansion and immune dysregulation.

### Compartment-Resolved Microbiome Analysis Reveals Low Stromal Biomass in Peri-Implantitis

Peri-implantitis has long been recognized as having a distinct microbial profile compared to periodontitis^27^; however, it remains unknown whether its rapidly advancing phenotype is attributable to dysbiosis^28^. Additionally, we have recently observed tissue-level, cell type and spatial heterogeneity in microbial localization in periodontitis^8,19^. Since we observed increased vascular-like structures in peri-implantitis immune lesions (Figure 1, Supplementary Figure 1), we hypothesized that peri-implantitis stroma might exhibit a higher bacterial burden than periodontitis, potentially driving exacerbated host responses related to local immune activation and tissue remodeling. To address this, we first used tissues sections for RNA fluorescence *in situ* hybridization (FISH), targeting bacterial *16S* rRNA to quantify microbial burden directly within FFPE tissue sections (Figure 2a). Unexpectedly, we observed that periodontitis samples consistently exhibited higher *16S* counts per cell across both epithelial and stromal compartments compared to peri-implantitis (Figure 2b,c).

**Figure 2.**
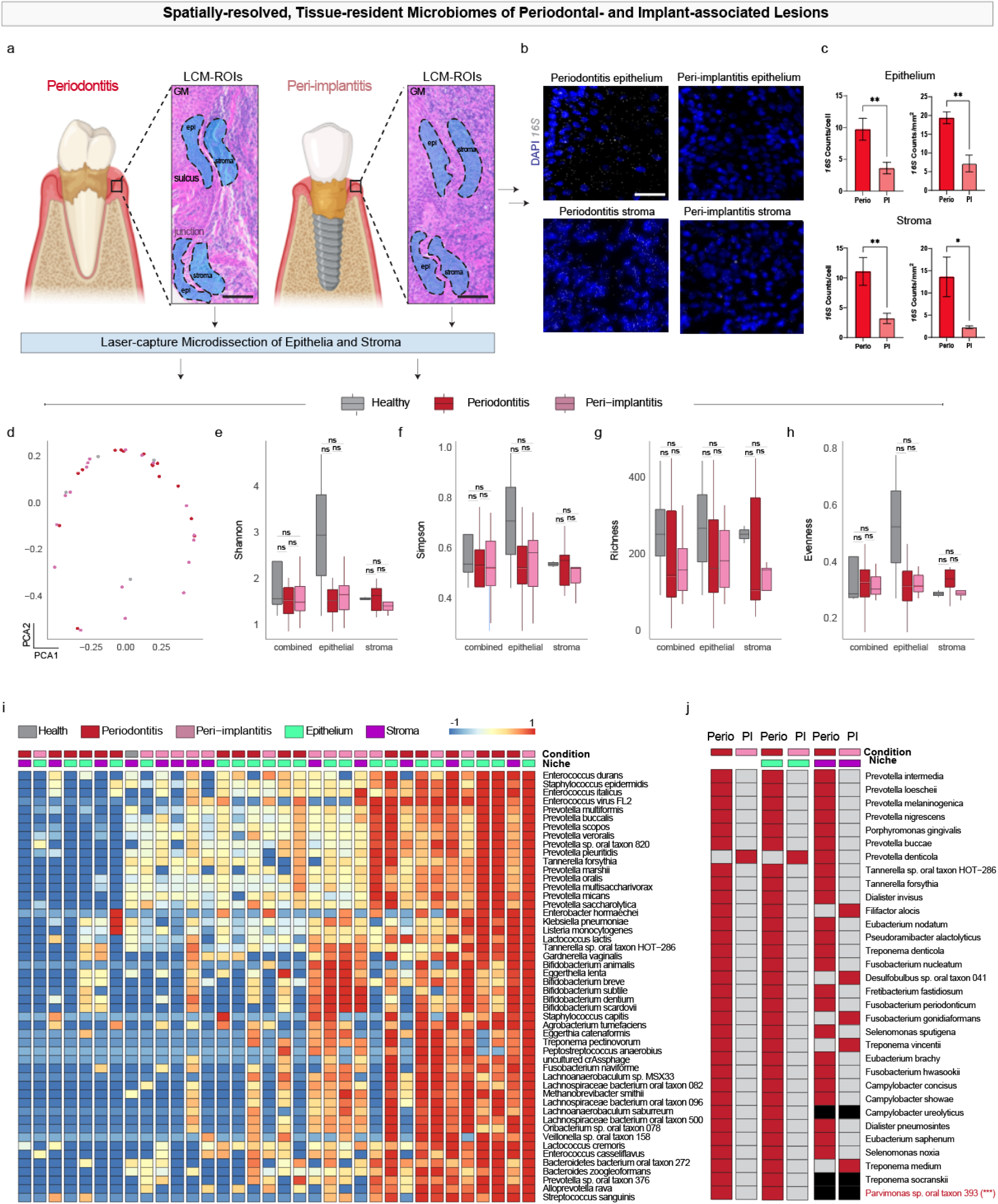
Dybiosis is not a major contributor to peri-implantitis pathogenesis. a) Using a probe for *16S* rRNA, we observed lower enrichment of bacterial signal in peri-implantitis epithelia and stroma. b) H&E images showing areas from which laser capture microdissection (LCM) samples were acquired. c)-g) represent first-in-class whole-genome sequencing-derived data of FFPE-derived LCM sections. c) We found few unique represented species comparing periodontitis and peri-implantitis. d) Alpha-beta diversity of represented species across health and diseases showed no significant species. e) Shannon and Simpson diversity indices, including richness and evenness across health and diseases, showed that while health differed from both periodontitis and peri-implantitis, in general, peri-implantitis had less diversity. f) Top represented species across health, periodontitis, and peri-implantitis samples. g) Directly comparing periodontitis and peri-implantitis for top-represented pathogens, only one species (*Parvimonas spp.*) was enriched in periodontitis; no other species statistically differed between the diseases. Scale bar: b) 20 µm For g) p < 0.0005. Images in a) were generated using BioRender.

To understand the diversity of the samples, we considered Shannon diversity to account for rare bacterial species, Simpson diversity to emphasize dominant species, richness to understand the number of distinct features, and evenness to determine how evenly individual counts were distributed among taxa (Figure 2d-h). We compared each of these four metrics across health, periodontitis, and peri-implantitis, considering combined compartments (epithelial- and stromal-specific compartments, Supplementary Figure 2). Across these indices, we found that in the combined data, there were comparatively few outliers, which resulted in no significant changes across any of the metrics. The widest variations occurred in healthy epithelia; this data is consistent with known microbiome composition differences in healthy individuals. We observed that while periodontitis and peri-implantitis epithelia had few differences, peri-implantitis stroma consistently had narrower indices than periodontitis stroma.

We then plotted the top enriched species, displaying compartments from least to most enriched (left to right; Figure 2i). No strong pattern emerged between diseases. We next extracted the top significant species across all 3 disease sets for comparison (Figure 2k). While we identified *Tannerella forsythia* as a top-represented species, we found few others that have been directly identified as contributing to pathogenesis. We reanalyzed the data for top contributing species using known periopathogens and asked whether any were enriched in periodontitis compared to peri-implantitis. While nearly all known periopathogens showed higher abundance in periodontitis, only *Parvimonas sp. oral taxon 393* was statistically significant; the trend matched for epithelia. In stroma, a difference was noted for enriched *Filifactor alocis*, *Treponema medium*, and *Treponema vincentii*^29^. Thus, the accelerated tissue destruction seen in peri-implantitis likely did not stem from dysbiosis alone.

### Vascular Endothelial Cell Dysfunction Emerges as a Core Pathogenic Feature in Peri-implantitis

Peri-implantitis lesions have been reported to display prominent vascular density, often juxtaposed to immune cell infiltrates^24^; this is a pattern observable in our own histological datasets (Figure 1). This raised the possibility that vascular remodeling may be a core and underappreciated contributor to disease pathogenesis, distinct from classical immune-driven tissue destruction. To investigate this, we integrated scRNA-seq datasets from peri-implantitis patient biopsies with the v1 Human Periodontal Atlas (HPA), generating an updated composite reference that encompasses health, gingivitis, periodontitis, and peri-implantitis (Figure 3a; proportions, Supplementary Figure 3a). Integration using Harmony enabled improved alignment across datasets and was superior to Seurat-based methods for batch correction (Supplementary Figure 3b, c).

**Figure 3.**
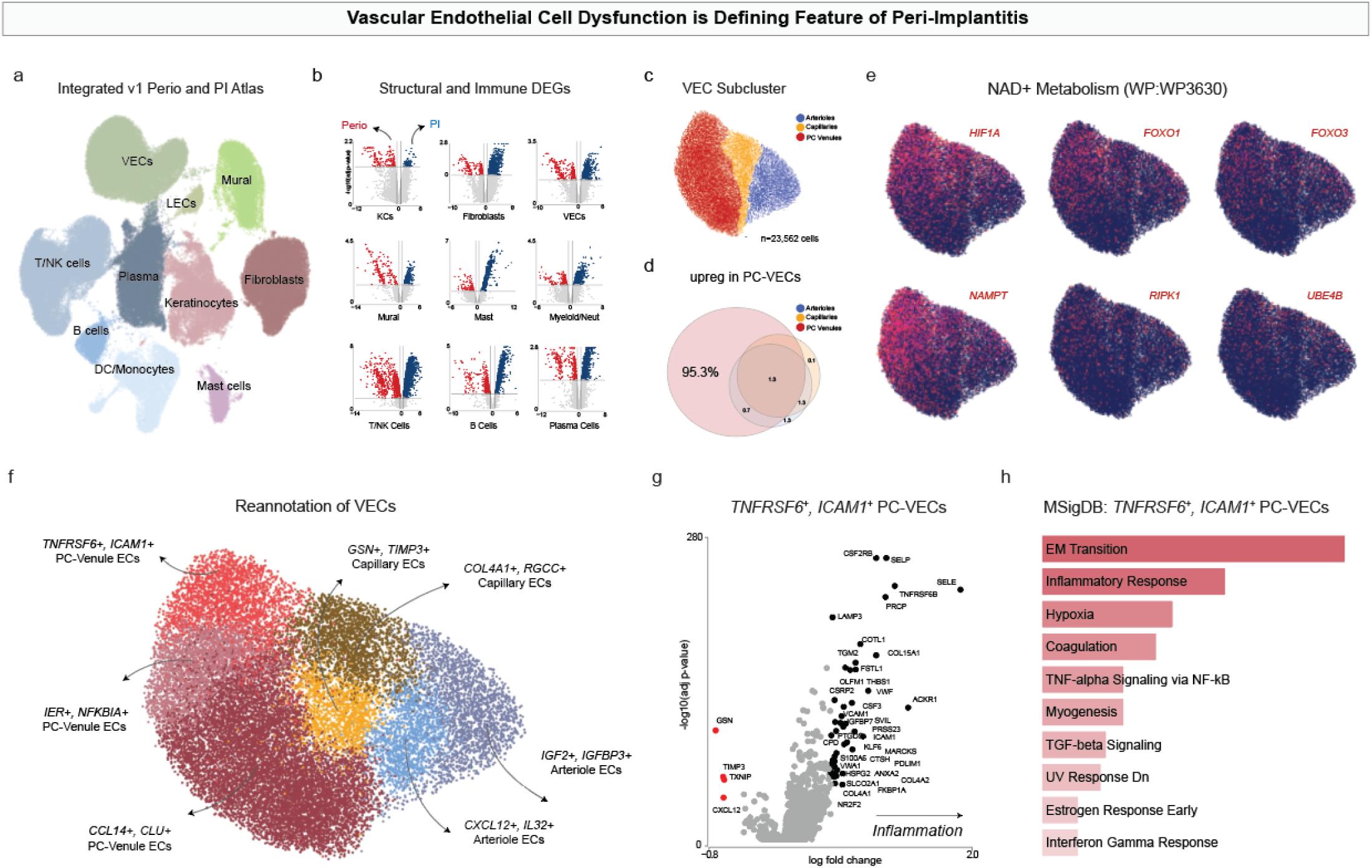
**Integration of single-cell RNA sequencing (scRNAseq) data reveals shared vascular populations across health and disease**. a) Our Human Periodontal Atlas (HPA) v1. b) Integration of peri-implantitis scRNAseq data resulted in our HPA v1.1. c) Focusing on the vasculature, we identified arterioles (A-VECs), capillaries (C-VECs), and post-capillary venules (PC-VECs). d) PC-VECs expressed more unique genes in peri-implantitis. e) differentially expressed gene analysis of arterioles, capillaries, and post-capillary venules revealed NAD^+^-associated metabolism in PCVs. f) Reannotation of VECs revealed among 8 subpopulations, a subpopulation of *TNFRSF6^+^*/*ICAM1^+^* PC-VECs expressed the NAD^+^-associated metabolism genes in e). g) Differentially expressed genes (DEGs) of *TNFRSF6^+^*/*ICAM1^+^*PC-VECs. h) Classifying the DEGs using MSigDB revealed inflammatory response, hypoxia, and TNF-alpha signaling as top represented pathways.

We then performed pseudobulk analysis of every major cell cluster after signature analysis (Supplementary Figure 3d), comprised of structural and immune cell types (Figure 3b; Supplementary Table 2). We performed differentially expressed gene (DEGs) analysis, and while there were many interesting trends upregulated in peri-implantits cell types compared to periodontitis, we focused on vascular endothelial cells (VECs) in peri-implantitis. Using this DEG list, pathway enrichment analysis revealed that peri-implantitis VECs exhibit a profound shift in both signaling and metabolic programs. We identified significant enrichment for growth factor signaling (EGFR and KIT receptor pathways), hormonal regulation (insulin, leptin, and thyroid hormone signaling), and proinflammatory responses (IL-6, TROP2, TNF-α via NF-κB; Supplementary Data 2). Notably, NAD^+^ metabolism and sirtuin signaling was selectively enriched in peri-implantitis VECs (Figure 3e, h).

The selective enrichment of NAD^+^ metabolism and sirtuin signaling in peri-implantitis VECs suggests that they are not only activated but metabolically stressed, implicating energy depletion and impaired repair capacity as key contributors to the accelerated breakdown observed in peri-implantitis. This was confirmed by the significantly downregulated pathways in peri-implantitis VECs, including oxidative phosphorylation (OXPHOS), mitochondrial complex I and IV assembly, and electron transport chain activity (Supplementary Data 2). Suppression of these programs, alongside the upregulation of inflammation, hypoxia, and endothelial remodeling pathways, suggests a shift from quiescent vascular maintenance to a dysfunctional, inflamed, and energetically compromised state.

From this integrated dataset, we extracted the VEC compartment and performed high-resolution subclustering, resolving three canonical VEC subtypes: arteriolar (A-VECs), capillary (C-VECs), and post-capillary venular vascular endothelial cells (PCV-VECs) as we have previously done (Figure 3c)^30^. Marker gene analysis confirmed their identity: arterioles expressed *SAT1, CLDN5, SRGN*, and *CXCL12*; capillaries expressed *COL4A1 and PVLAP*; and PC-VECs expressed *SELP, SELE*, and *ACKR1* (Supplementary Figure 3e). DEG analysis comparing each subpopulation between peri-implantitis and periodontitis revealed that peri-implantitis-PCV-VECs were greatly affected, upregulating >1000 genes that were unique when compared to what was upregulated in A-VECs and C-VECs, which only displayed a small fraction of DEGs (Figure 3d). This pointed to PCV-VECs being a significant contributor to the endothelial cell dysfunction observed in peri-implantitis. When considering genes for hypoxia (*HIF1A*), NAD+ synthesis (*NAMPT*), and other genes related to NAD+ metabolism (*FOXO1, FOXO3, RIPK1, UBE4B*), we found that these were almost exclusively concentrated in a subregion of PCV-VECs.

To be better understand these cells, we performed a Tier 4 cell type analysis using clustered regions from Leiden clustering. We found two types of A-VECs (1: *CXCL12^+^; IL32^+^* and 2: *IGF2^+^, IGFBP3^+^*), two types of C-VECs (1: *GSN^+^, TIMP3^+^* and 2: *COL4A1^+^ and RGCC^+^*) and three types of PCV-VECs (1: *CCL14^+^, CLU^+^;* 2: *IER^+^, NFKBIA^+^;* and 3: *TNFSFR6^+^, ICAM1^+^*). It was these latter PCV-VECs that aligned with the NAD+ metabolism (Supplementary Table 2). We then repeated DEG analysis of these *TNFSFR6^+^*/*ICAM1^+^*PCV-VECs, finding in inflammation compared to health many upregulated genes within this cluster (Figure 3g). Pathway enrichment analysis of *TNFRSF6B^+^*/*ICAM1^+^* PCV-VECs revealed a variety of disease-relevant programs, including endothelial-to-mesenchymal transition, TNF-α/NF-κB signaling, TGF-β signaling, hypoxia, and IFN-related inflammatory responses (Figure 3h). These signatures implicate PCV-VECs not only as passive conduits but as active participants in immunovascular signaling, potentially sustaining chronic inflammation in peri-implant lesions. Unlike traditional models of periodontal disease centered on immune cells or microbial dysbiosis, these data suggest that vascular cells themselves may initiate or perpetuate tissue pathology.

### Endothelial Remodeling and CD38⁺ Vascular Expansion Define Peri-Implantitis Immune Niches

Having discovered VECs as important cell types for peri-implantitis pathogenesis, we decided to investigate the spatial immune and stromal architecture underlying peri-implantitis pathogenesis using sp-proteomics. Using healthy and periodontitis samples from our HPA v1.1^8^, we performed the Phenocycler-Fusion assay (Akoya Biosciences) on peri-implantitis tissues (Figure 4a). Our 32-antibody CODEX panel targeted lineage, state, and activation markers for epithelial, stromal, and immune compartments, including VECs (Figure 4b). Following acquisition, we implemented a deep-learning segmentation model trained on multi-input features, including DAPI and membrane markers, enabling accurate single-cell segmentation across complex inflammatory niches (Figure 4c).

**Figure 4.**
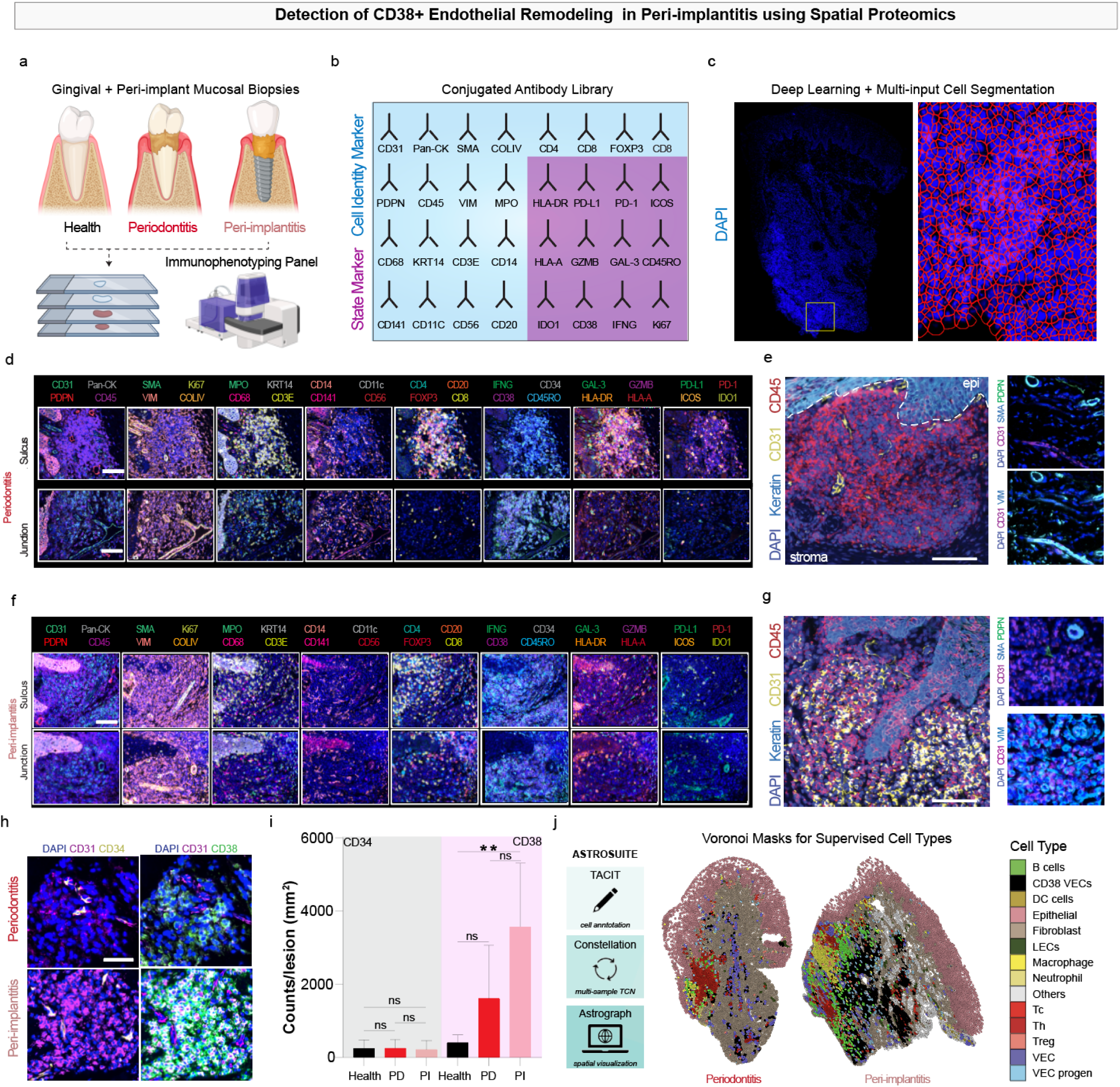
Highly multiplexed immunofluorescence unveils the immune heterogeneity in periodontitis and peri-implantitis. a) Schematic of immunophenotyping of health, periodontitis, and peri-implantitis. b) Conjugated antibody library used in this study to distinguish cell identity and state. c) We trained CellPose for cell segmentation based on DAPI to enable precise fluorescence calls. d)-g) Regional immunophenotypes of the junction and sulcus in d) periodontitis were defined by e) immune expansion, yet in f) peri-implantitis, g) CD31^+^/VIM^+^ vascular cells expanded without additional identifying markers. h)-i) CD31^+^/CD34^+^ progenitor cells were present in both tissue types, yet we observed expansion of CD31^+^/CD38^+^ VECs in peri-implantitis lesions. j) Using our AstroSuite pipeline, we used the segmented data to generate Voronoi masks of supervised, defined cell types. Scale bars: d)-g): 50 µm; h): 20 µm. Statistical analysis: i) **p < 0.005. Images in a and j were generated using BioRender.

We then began generating cell recipes using the structural and immune markers in periodontitis and peri-implantitis (Figure 4d; health, Supplementary Figure 4). PanCK^+^/KRT14^+^ marked epithelial cells, whereas stromal cells were marked by vimentin (VIM^+^); Ki67^+^ identified cycling and non-cycling cells within each. We initially refined endothelial cells by classifying CD31^+^ cells into VECs, VEC progenitor (CD34^+^), and lymphatic endothelial cells (LECs; PDPN^+^). Using CD45^+^ as the overarching immune cell marker, in combination with antigen-presenting (HLA-A^+^/HLA-DR^+^), we classified adaptive CD3E^+^ T cells and CD20^+^ B cells and innate CD56^+^ NK cells, CD11c^+^/CD141^+^ dendritic cells, myeloperoxidase (MPO^+^) neutrophils, and CD68^+^ macrophages. T cells were further refined into cytotoxic (CD8^+^), helper (CD4^+^), and regulatory (CD4+/FOXP3^+^) T cell types.

Initial inspection of pan-CK^+^ epithelium and surrounding VIM^+^ stroma revealed significant expansion of disorganized, CD31^+^ vasculature in peri-implantitis relative to periodontitis (Figure 4d-g). VECs exhibited partial loss of endothelial identity and increased expression of mesenchymal markers (e.g., VIM) without concomitant expression of other vasculature markers (e.g., PDPN, SMA), consistent with an EndoMT-like phenotype that may exacerbate tissue fibrosis and immune cell recruitment. This prompted focused evaluation of VECs and their phenotypic diversity. CD31^+^ VECs in peri-implantitis lesions exhibited frequent co-expression of CD38, a marker associated with inflammation and NAD^+^ metabolism in immune^31^ and endothelial cells^32^. CD38^+^ VECs were CD20-negative, and absent or rare in healthy or periodontitis tissues (Figure 4d–g). Importantly, this expansion was not paralleled by a rise in CD34^+^ VEC progenitors per area (Figure 4h,i), suggesting that CD38^+^ endothelial remodeling reflects an inducible, disease-specific cell state rather than a developmental program.

To further characterize the immune-stromal niches enriched in peri-implantitis, we deployed the *AstroSuite* pipeline (*TACIT*, *Constellation*, and *Astrograph*)^7,20^, which integrates spatial cell annotations, tissue compartment mapping, and spatial multiomics analyses (see Supplementary Information). Using Voronoi-based neighborhood reconstruction, we resolved spatial relationships among immune cells (CD8^+^ T cells, macrophages, Tregs), structural elements (keratinocytes, LECs, fibroblasts), and VEC subsets. On a qualitative basis, as expected, the peri-implantitis immune lesion was dramatically larger than the periodontitis lesion (quantitative; Figure 1g); however, this also revealed an expansion of CD38^+^ VECs in peri-implantitis, displaying multiple perivascular immune niches and vascular islands that were found to be distal to the originating lesion in peri-implantitis, (Figure 4j). These findings identify CD38^+^ endothelial remodeling as a hallmark of peri-implantitis and highlight the utility of spatial proteomics and *AstroSuite* in resolving complex vascular-immune niches.

### CD38⁺ Endothelial Cell Niches Distinguish the Spatial Architecture of Peri-Implantitis

We next asked how CD38^+^ vascular endothelial cells (VECs) were spatially organized in peri-implantitis and whether they co-occurred with specific immune or stromal populations (Figure 5a). To explore this, we included CD31^+^/CD38^+^ VECs as a distinct cell identity in our *TACIT*-based pipeline. Triwise enrichment analysis comparing *TACIT*-assigned cell types across health, periodontitis, and peri-implantitis revealed that CD38^+^ VECs, CD31^+^/CD34^+^ VEC progenitors, and CD31^+^/CD38^⁻^ classical VECs were most enriched in peri-implantitis (Figure 5b). In contrast, immune cells, such as B cells, helper T cells, macrophages, and neutrophils, were more concentrated in the lesions of both diseases. Periodontitis samples were more relatively enriched for NK cells, cytotoxic T cells, and dendritic cells. Both disease states also showed a relative depletion of epithelial and regulatory T cells compared to healthy tissue. These patterns were confirmed in lesion-wide bar plots, highlighting CD38^+^ VECs as the most defining cell population in peri-implantitis (Figure 5c). Further, when we analyzed immune and vascular activation, CD38^+^ VECs were frequently enriched for IFNγ (Supplementary Figure 5).

**Figure 5.**
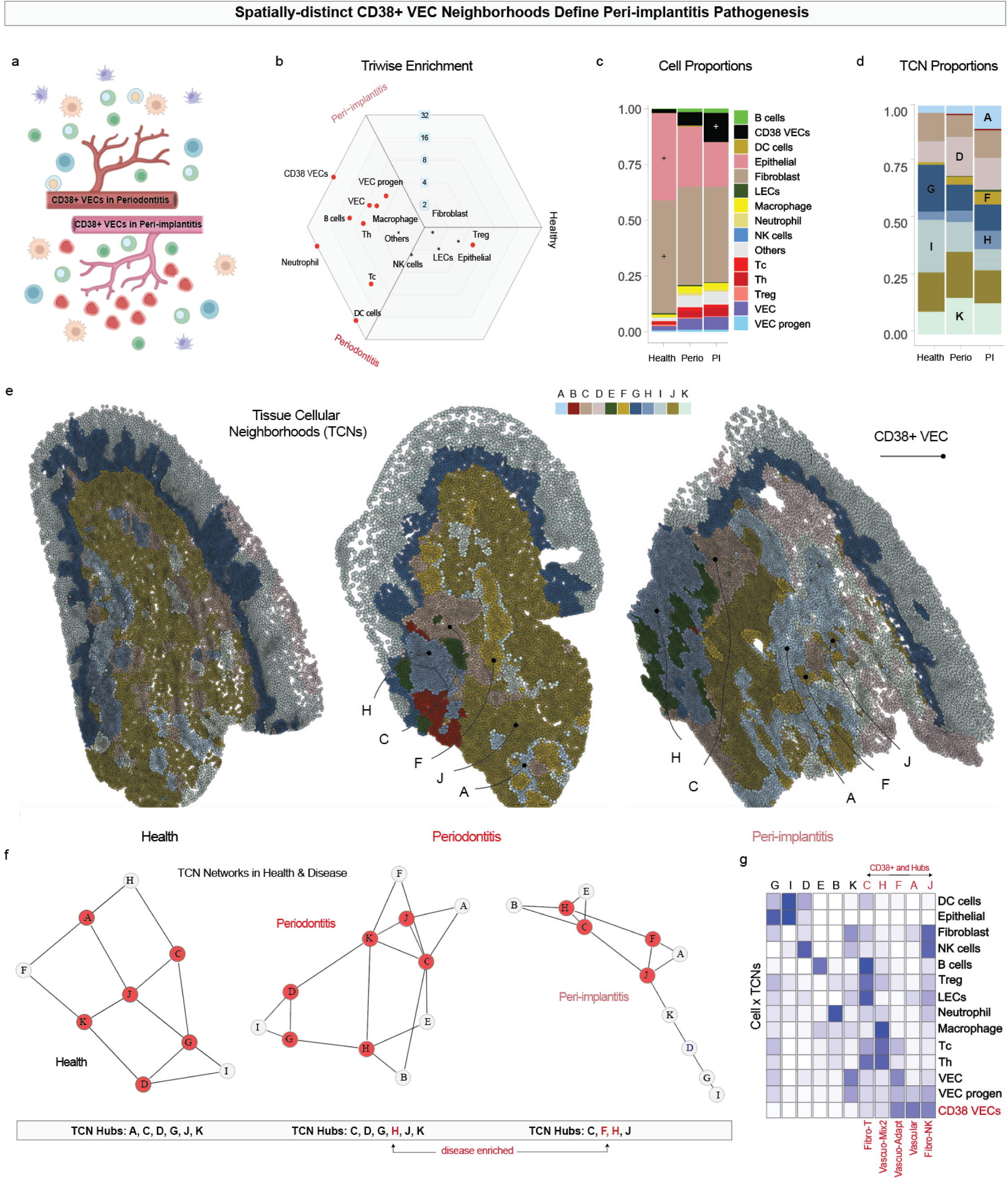
Peri-implantitis tissue-connected neighborhood proportions and spatial localizations define a unique pathology dominated by CD38 VECs. a) Schematic of different cell environments surrounding CD38^+^ VECs in periodontitis and peri-implantitis. b) Triwise enrichment comparison of health, periodontitis and peri-implantitis revealed the periodontitis/peri-implantitis group was enriched for most immune cell types, yet peri-implantitis was strongly enriched for CD38^+^ VECs. c) Proportions of cells across health and disease states. d) Proportions of TCNs within health and disease states. e) Voronoi representations of healthy, periodontitis, and peri-implantitis tissue connected neighborhoods (TCNs). Peri-implantitis shows expansion of CD38^+^ VECs through TCNs A, F, and H. f) TCN interaction hubs are dominated by CD38^+^ VEC neighborhoods in peri-implantitis. g) Analysis of Cell x TCN information revealed five hubs enriched for CD38^+^ VECs; TCNs A, F, and H all expand in peri-implantitis. Images in a) were generated using BioRender.

To analyze the spatial architecture of peri-implantitis, we used *AstroSuite*’s *Constellation* tool to assign cells to 11 tissue-connected neighborhoods (TCNs) based on co-occurrence and proximity (Figure 5d). Healthy samples were enriched for TCNs G and I; periodontitis for TCNs D and K; and peri-implantitis for TCNs A, F, and H (Figure 5e). CD38^+^ VECs were prominently represented in TCNs A, F, and J: TCN A was an exclusion zone composed nearly entirely of CD38^+^ VECs; TCN F showed CD38^+^ VECs intercalating with classical VECs, VEC progenitors, and cytotoxic T cells; and TCN J consisted of CD38^+^ VECs adjacent to fibroblasts. These findings revealed two distinct spatial modes for CD38^+^ VECs in peri-implantitis: self-contained vascular niches and intercalated vascular-immune niches.

Next, we evaluated how these neighborhoods restructured tissue-wide architecture (Figure 5f). In health, a balanced network of six interconnected TCN hubs (A, C, D, G, J, K) with three or more connections to other TCNs supported tissue homeostasis. Analyzed periodontitis tissues apparently lost a “homeostatic” hub A and gained pathologic one, hub H, which we found is a new immune-rich stromal hub near the lesion front containing macrophages, T cells, and VECs. In peri-implantitis, TCNs A, D, G, and K were lost as hubs and replaced by TCNs H and F. TCN F formed a novel vascular-immune hub dominated by CD38^+^ and classical VECs and cytotoxic T cells, indicating that vasculature was central to tissue remodeling in peri-implantitis.

To further characterize each TCN, we examined cell composition (Figure 5g). TCN A was dominated by CD38^+^ VECs and VEC progenitors; TCN F contained classical and CD38^+^ VECs, VEC progenitors, and cytotoxic T cells; and TCN J included fibroblasts, LECs, and NK cells alongside CD38^+^ VECs. TCN B and E were enriched for neutrophils and B cells, respectively. TCN C contained a mix of stromal and immune cells, such as B cells, helper and regulatory T cells, and LECs, suggesting it represented an ectopic lymphoid structure. TCN H was enriched for macrophages and all T cell subtypes, whereas TCNs G and K marked transitional and structural stroma, respectively. These data reveal that CD38^+^ VECs in peri-implantitis are embedded within structurally distinct spatial neighborhoods that alter hub connectivity and stromal-immune architecture. Rather than simply increasing in number, CD38^+^ VECs reorganize the lesion’s spatial framework, establishing exclusion and intercalation zones that shape the immune landscape of chronic disease. Furthermore, this recurring alignment between CD38^+^ VECs and cytotoxic T cells within TCN F suggests a specialized vascular-immune niche that may coordinate immune effector positioning and tissue remodeling in peri-implantitis, highlighting a spatially orchestrated vascular expansion as well as vasculopathy.

### Convergent CD38^+^ Endothelial Pathology in Rapidly advancing Periodontitis and Peri-implantitis

To test whether the CD38^+^ vasculopathy and stromal-immune remodeling observed in peri-implantitis are unique to the implant microenvironment or instead represent a broader pathological motif in destructive oral inflammatory diseases, we examined tissues from patients with Stage III, Grade C periodontitis. This rapidly advancing form of periodontitis is characterized by rapid tissue breakdown, even in systemically healthy individuals, offering an opportunity to assess whether similar spatial vascular-immune modules exist in natural-tooth disease. We collected matched gingival biopsies from unaffected and affected sites within the same individual (Figure 6a) and first evaluated tissue architecture using hematoxylin and eosin (H&E) staining. Compared to unaffected regions, affected sites exhibited clear histopathological features of disease, including stromal expansion, dense leukocyte infiltration, and epithelial rete ridge broadening (Figure 6b). These histological signatures resembled those observed in peri-implantitis and prompted spatial proteomic profiling using the sp-proteomics (Akoya PhenoCycler Fusion) panel.

**Figure 6.**
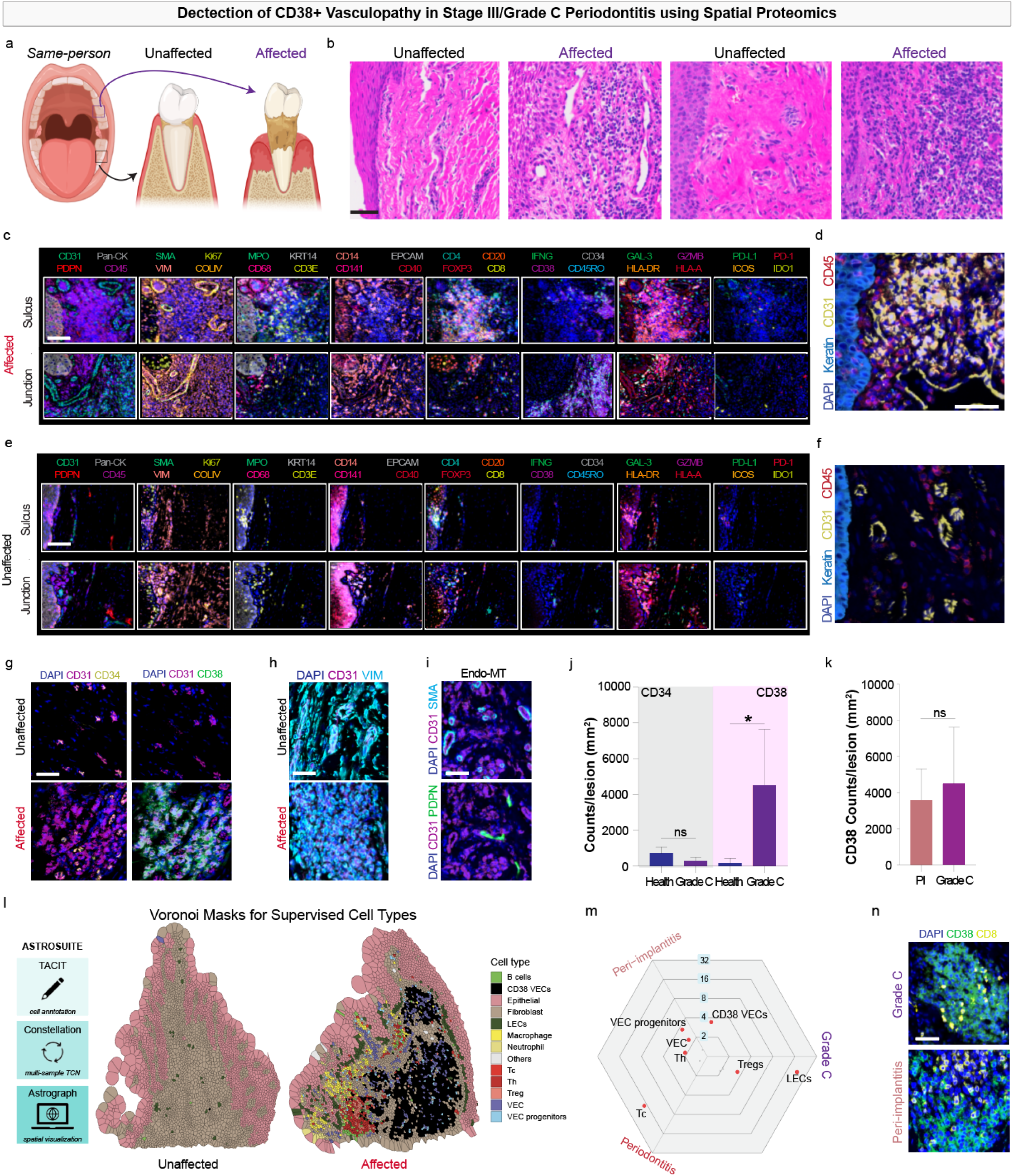
CD38^+^ vasculopathy is a feature of Stage III/Grade C periodontitis. a) Schematic showing same-person collection of unaffected and affected tissues. Triwise plot showing enriched cell types shared between different tissue states. b) H&E analysis shows affected sites display significantly higher cellular density. c)-f) Regional immunophenotypes of affected and unaffected sites show that c) Grade C periodontitis lesions display immune infiltrant and d) CD31^+^ expansion, while e),f) unaffected sites appear comparatively healthy. g) Inspection of the expanding CD31^+^ lesions revealed CD38^+^, not CD34^+^ VEC enrichment. h),i) We further observed these cells were h) CD31^+^/VIM^+^ in affected sites and i) were not enriched for other vascular identifying markers. j) Grade C periodontitis sites were enriched for CD31^+^/CD38^+^ VECs. k) We observed no differences in lesion density of CD31^+^/CD38^+^ VECs in peri-implantitis and Grade C periodontitis. l) We applied our AstroSuite toolkit to unaffected and unaffected sites and observed tissue-wide expansion of CD38^+^ VECs. m) Triwise analysis of three disease types revealed peri-implantitis and Grade C periodontitis were enriched for CD38^+^ VECs. n) Both diseases shared the predicted CD38^+^ VEC/CD8 T cell neighborhood predicted by Constellation. Scale bars: c)-f): 50 µm; g)-i), n): 20 µm. Statistical analysis: j) *p < 0.05. Images in a and l were generated using BioRender.

High-plex imaging revealed consistent expansion of CD31^+^/CD38^+^ VECs in affected Grade C tissues, localized prominently in connective tissue regions beneath the sulcus and junctional epithelium (Figure 6c). These CD38^+^ VECs frequently appeared as organized vascular clusters co-localizing with CD45^+^ leukocytes and basal epithelial zones (Figure 6d), forming similar patterns of vascular remodeling observed in peri-implantitis yet not in matched healthy sites (Figure 6e,f). To probe for features of endothelial plasticity similar to peri-implantitis, we assessed co-expression of CD38 with stromal markers. We observed upregulation of vimentin (VIM) in CD38^+^ VECs in affected—but not unaffected—tissue regions, without upregulation of PDPN or SMA and lack of expansion of CD34, consistent with endothelial-to-mesenchymal transition (EndoMT) (Figure 6g–j). Additionally, we observed similar counts of CD38 per lesion measurement area when compared to peri-implantitis (Figure 6k), suggesting that CD38^+^ vasculature in Grade C lesions may also participate in immune guidance and undergo structural remodeling.

To understand the cell distribution within tissues, we applied our *AstroSuite* pipeline to both the healthy and affected sites (Figure 6l). In affected tissues, we observed an expanding CD38^+^ VEC front similar to peri-implantitis. To uncover any shared features, we employed our triwise enrichment analysis of *TACIT*-assigned cell types to compare low-grade (B) periodontitis, peri-implantitis, and high-grade (C) periodontitis (Figure 6m). We found that across all three disease states, only CD38^+^ VECs were shared between peri-implantitis and Grade C periodontitis. Further, spatial mapping revealed frequent co-localization of CD38^+^ VECs and CD8^+^ cytotoxic T cells within the same tissue niches (Figure 6n). These immune-vascular neighborhoods were repeatedly observed in affected sites and were notably absent in unaffected tissues. This mirrors the peri-implantitis phenotype, where CD38^+^ vasculature interfaces directly with immune effectors, specifically CD8+ cytotoxic T cells. The recurrence of these patterns across disease types suggests a conserved stromal-immune architecture underpinning destructive inflammation in the oral cavity. Overall, these data extend our findings beyond peri-implantitis and support a model in which CD38^+^ VEC expansion, EndoMT, and selective immune cell coupling form a shared spatial program in rapidly progressive inflammatory lesions.

## DISCUSSION

The oral cavity presents a powerful yet underutilized model for investigating tissue remodeling in human disease. Here, mechanical stress, polymicrobial exposure, and continuous immune surveillance converge within an anatomically confined, readily accessible tissue. Unlike many mucosal surfaces, the oral mucosa exhibits both rapid and spatially restricted inflammatory dynamics, offering a rare opportunity to observe lesion development in real time—from acute shifts over days to long-term remodeling over months or years. Leveraging these advantages, we conducted what we believe to be the first spatial omics study of both peri-implantitis and Stage III/Grade C periodontitis. This integrated multi-disease spatial analysis, which we believe is among the first of its kind in the spatial biology era, uncovered a shared immunovascular pathology defined by the expansion of CD38^+^ vascular endothelial cells. These cells do not merely reflect downstream inflammation but actively reshape the tissue landscape by forming discrete, lesion-specific spatial modules that couple with cytotoxic T cells and disrupt classical fibrovascular architecture.

A review of the literature underscores that endothelial cell biology remains understudied in oral inflammatory diseases. Most of the limited spatial or single-cell omics studies in the field have focused on immune, fibroblast, and/or epithelial dynamics, while vasculature is often treated as a static scaffold. Yet endothelial dysfunction plays a critical role in many chronic pathologies, where aberrant endothelial states reshape immune infiltration and tissue-specific immunity^33,34^. By anchoring our analysis around endothelial subtypes, we reveal that CD38^+^ VECs emerge consistently in rapidly advancing oral lesions, co-occur with both innate and adaptive immune cells, and show signatures of IFNγ exposure and endothelial-to-mesenchymal transition (Supplementary Figure 5). These cells are not only expanded but spatially reorganized, forming distinct exclusion or intercalation zones across tissue compartments.

Between the two rapidly advancing diseases studied here, peri-implantitis is rising in prevalence alongside increased dental implant use, affecting roughly one-quarter of patients^25^; for a variety of factors, the prevalence of peri-implantitis is expected to increase^35^. Peri-implantitis is often characterized as a biofilm-driven inflammatory lesion, yet our findings revise this bacteria-centric view by revealing a consistent expansion of CD38^+^ VECs and immune-coupled vascular hubs, despite relatively lower bacterial abundance in some high-grade cases. Importantly, the current therapeutic paradigms for peri-implantitis center on plaque control and debridement but have yielded limited success in halting disease progression^36^. We hypothesize that part of that limited success is because the host-response is uniquely shaping niche-specific tissue breakdown, which may be exacerbated but not solely driven by dysbiosis. Similarly, in Stage III/Grade C periodontitis, we observe parallel expansion of CD38^+^ VECs and their association with cytotoxic T cells in the connective tissues. This form of periodontitis is also difficult to treat^16^, a general feature of oral inflammation based on our current studies outside this niche.

This vascular-centric spatial phenotype introduces a new conceptual framework for understanding chronic oral disease and suggests novel therapeutic avenues. CD38 is an NAD^+^-hydrolyzing ectoenzyme that also serves as an immune modulator. Its expression on endothelial cells has been implicated in inflammatory vasculopathies of the heart, brain, and kidneys^37–39^. Monoclonal antibodies against CD38 (e.g., daratumumab, isatuximab) are already FDA-approved for hematologic malignancies and could be repurposed, using local delivery strategies, to target pathogenic endothelial states in mucosal inflammation as they have been recently explored for autoimmunity^40–42^. Moreover, interventions aimed at restoring NAD^+^ levels, activating SIRT1, or blocking IFNγ signaling, pathways observed in the CD38^+^ VECs in our data, may blunt maladaptive vascular remodeling and its downstream immune effects. These targets provide a translational bridge between spatial pathology and molecular precision therapeutics in oral medicine.

The diagnostic implications of our findings are equally significant. Traditional metrics for lesion severity focus on immune composition, microbial load, or tissue damage^43^. Yet our analysis shows that endothelial subtypes and their spatial arrangements better predict tissue organization and immune residency than immune proportions and stromal density alone. For instance, CD8^+^ T cells co-occur with CD38^+^ VECs in peri-implantitis and Grade C periodontitis but not in classical periodontitis, despite CD8^+^ T cells and CD38^+^ VECs being present in all three diseases. This indicates that spatial context, not just cell identity, determines immunopathology. Future diagnostic frameworks may incorporate vascular profiling to stratify disease subtypes and predict treatment response, a strategy applicable not only in the oral cavity but across mucosal diseases and disorders.

Despite its strengths, our study has limitations. The spatial omics analyses were performed on a carefully curated yet limited set of human tissue samples, reflecting the difficulty in acquiring high-quality peri-implant and Stage III/Grade C periodontitis lesions. While the consistency of CD38^+^ VEC emergence across samples is striking, larger cohorts are needed to enable subtyping and longitudinal tracking. Additionally, the lack of healthy peri-implant mucosa or peri-implant mucositis samples constrains our ability to resolve early transitions in vascular remodeling. Finally, current animal models lack fidelity for replicating human-specific endothelial–immune interactions and NAD^+^ metabolism from our experience, complicating preclinical translation.

Nonetheless, this study reframes vascular architecture as a central player in oral tissue inflammation and destruction. The emergence of CD38^+^ VECs within two distinct diseases points to a shared vascular module that disrupts immune equilibrium and structural integrity.

These lesions are not merely consequences of immune overactivation or bacterial invasion, but spatially organized outcomes of endothelial rewiring. As spatial biology becomes increasingly integrated into translational science, we propose that vascular-centric frameworks may yield new insights across organ systems. The oral cavity, with its accessibility and spatiotemporal resolution, provides a powerful setting to pioneer these approaches. In our view, this spatial omics study lays the foundation for incorporating vascular profiling into the next generation of oral disease classification and treatment strategies, and potentially into broader models of mucosal pathobiology^44^.

## MATERIALS AND METHODS

### Ethics statement

This research complies with all relevant ethical regulations. All studies using human gingival biopsies were approved by either the University of Pennsylvania (IRB#6; Protocol #844933; Lead PI: KIK), the National Institutes of Health through the National Institute of Dental, Oral and Craniofacial Research (NCT#01805869; Lead PI: Janice Lee [NIDCR]) or the University of Campinas FOP-UNICAMP Research Ethics Committee (51043921.6.0000.5418; Lead PI: RC).

### Single-cell analyses of RNA sequencing data

Integrated periodontitis + peri-implantitis single-cell RNA sequencing (scRNAseq) atlas generation and subclustering using Trailmaker. We have previously generated an integrated scRNAseq atlas of human periodontium, including health, gingivitis, and periodontitis samples as previously described^8^. Biopsies acquired from patients with peri-implantitis were digested, stained, and sorted according to this previous protocol. We adapted the same parameters to ensure consistent data treatment and accurate integration of the peri-implantitis single-cell data. In brief, we created a copy of our Trailmaker project and uploaded pre-filtered count matrices for peri-implantitis data to the project. We used the following parameters: 1) lower than 500-UMI barcodes were filtered out; 2) mitochondrial reads with >15% indicated dead and dying cells and were filtered out; 3) outliers were filtered using the MASS package, which removed droplets outside lower and upper boundaries; and 4) barcodes with doublet scores greater than 0.5 were filtered out. The remaining samples, containing 300-8000 high-quality barcodes, were integrated. We used the Harmony R package for batch correction and Seurat’s implementation of the Louvain method for clustering^45^. Uniform Manifold Approximation and Projection (UMAP) was performed using Seurat’s wrapper around the UMAP package^46^. Cluster-specific marker genes were identified using the presto package implementation of the Wilcoxon rank-sum test. We subclustered vascular endothelial cells (VECs) by classifying cells according to Tier 1 type, then subsetted them into their own Trailmaker project. We implemented the same integration pipeline as the full experiment and annotated cells manually using the available literature and CellTypist^47^.

Transfer of Trailmaker data to CELLxGENE We exported our data and transferred it to CELLxGENE as previously described.

<u>Differentially expressed gene (DEG) analysis using Trailmaker and g:Profiler</u>. All cell subpopulations were grouped using the lasso tool on Trailmaker to enable pseudobulk RNA sequencing analyses at Tier 1 and, for VECs, Tier 2, Tier 3, and Tier 4 resolution. DEG lists and volcano plots were generated in Trailmaker, exported as .csv files, and uploaded to g:Profiler^48^. The .csv files are available in Supplementary Data 2 for further analysis.

### Spatial analyses conducted using gingival biopsies

<u>Tissue preparation, mounting, sectioning, and deparaffinization for mIF and RNA FISH</u>: Deidentified human gingival tissues were acquired from gingival biopsies (University of Pennsylvania, IRB #844933, MTA #68494; NIDCR, NCT#01805869; University of Campinas, #51043921.6.0000.5418). Peri-implantitis and Grade C periodontitis samples were selected according to previously established inclusion criteria^49^. Immediately after extraction, healthy and peri-implantitis tissues were placed in a 10% solution of N-buffered formalin (NBF) and fixed for 24 h in a 4°C refrigerator. After fixation, the tissues were washed twice in 1X PBS before being placed in 70% EtOH in a 4 °C refrigerator until they were ready to be mounted. Grade C periodontitis and matched tissues were similarly collected but were fixed at rt and were rinsed with tap water for 1 h before storing in EtOH prior to mounting. Tissues were embedded in paraffin blocks using a Leica system and stored in a 4 °C refrigerator until sectioning using RNAse precautions on a Leica system.

Formalin-fixed, paraffin-embedded (FFPE) human gingival tissue on SuperFrost Plus slides was heated to 60 °C on a slide warmer for 30 min. Following deparaffinization for 10 min using HistoChoice Clearing Agent (Electron Microscopy Services #64114-01), the tissues were rehydrated using a series of ethanol solutions (100%, 90%, 70%, 50%, and 30% EtOH in nuclease-free water) for 10 min (for 100%) and 5 min each, followed by 2 x 5 min in 100% nuclease-free water. During rehydration, 50 mL of a 1X solution of AR9 buffer (commercially available pH 9 buffer, Akoya Biosciences) in nuclease-free water was prepared and added to a Coplin jar. Following rehydration, the slides were added to the 1X AR9 buffer and covered with aluminum foil. Samples were antigen retrieved in a pressure cooker for 15 min at low pressure. Following antigen retrieval, the Coplin jar was removed from the pressure cooker and cooled for at least 30 min. The slide was then soaked in nuclease-free water for 30 s, followed by soaking in 100% EtOH for 3 min, both in Coplin jars.

<u>PhenoCycler-Fusion (PCF) on Human Tissues</u>: All reagents in this section were purchased and used as received from Akoya Biosciences unless otherwise noted. Samples underwent deparaffinization, rehydration, and antigen retrieval as described above. Following sample immersion in EtOH, the sample was immersed in Akoya Hydration Buffer for 2 min, followed by Akoya Staining Buffer for 20 min. While the sample cooled to rt, the antibody cocktail was prepared. To 362 µL of Akoya Staining Buffer was added 9.5 µL each of N, J, G, and S blockers. Then, 158 µL of blocking solution was pipetted into a 1.5 mL vial, and 1 µL of barcoded antibody (table below) was added to the vial such that the final volume of antibody diluent was 190 µL for each slide. After immersion in Staining Buffer, the slide was removed, the back and area around the sample were wiped dry, and the slide was added to a humidity chamber. *As a modification to the manufacturer’s instructions*, the antibody diluent was added to the sample, and the humidity chamber was placed in a 4°C refrigerator overnight. After removal of the blocking solution, the slide was placed in staining buffer for 2 min, followed by post-stain fixing solution (10% PFA in staining buffer) for 20 min. Following 3 x 2 min washing in 1X PBS, the slide was immersed in ice-cold MeOH for 5 min. While the slide was immersed, the final fixative solution was prepared by adding 1 vial of fixative (20 µL) and 1 µL of stock DAPI to 1 mL of PBS. The slide was removed from MeOH and placed in the humidity chamber, and 200 µL of the final fixative solution was added to the sample. This was left in place for 20 min. Then, the final fixative solution was removed, and the slide was washed in 3 x 2 min in 1X PBS.

To convert the slide into a flow cell for use in the PCF experiment, the back of an Akoya flow cell top was removed, and the top was placed adhesive face up in the Akoya-provided impressing device. The slide was removed from the 1X PBS, and the edges around the slide that matched where the top of flow cell adhesive would adhere were dried using a micro-squeegee toolkit (Essential Bangdi). Then, the slide which formed the bottom of the flow cell was placed sample-side down on the top of the flow cell without applying pressure to the adhesive. The tray of the impressing device was inserted into the device, and the lever was gently pulled to adhere to the top and bottom of the flow cell. After 30 s, the lever was depressed, the tray was pulled out, and the flow cell was removed. This flow cell was placed in 1X PCF buffer without buffer additive for a minimum of 10 min before any PCF experiment to allow for improved adhesion between the top and bottom of the flow cell.

To prepare the PCF reporter wells, a 15 mL Falcon tube was wrapped with aluminum foil. To this Falcon tube was added 6.1 mL of nuclease-free water, 675 µL 10X PCF buffer, 450 µL PCF assay reagent, and 4.5 µL of in-house prepared concentrated DAPI such that the final DAPI concentration was 1:1000. Then, this reporter stock solution was pipetted to 18 amber vials, with the volume in each vial being 235 µL. To each vial was added 5 µL of reporter per cycle. The total volume per vial was either 245 µL for a cycle with 2 reporters or 250 µL for a cycle with 3 reporters; to optimize reagents and reporters, no cycles contained only 1 reporter. Only one criterion was used to create a cycle: each cycle could contain a maximum 3 reporters, corresponding to 1 of Atto550, AlexaFluor 647, and AlexaFluor 750 (where appropriate; see below for more information). A separate pipet tip was used to pipet the contents of each amber vial to a 96-well plate. DAPI-containing vials were pipetted into a well in the H-row, whereas vials containing reporters were pipetted into wells in other rows. Once all wells were filled, Akoya-provided foil was used to seal the wells. Imaging was performed using a PhenoImager Fusion connected to a PhenoCycler i.e., the PhenoCycler Fusion 1.0 and 2.0 system (Akoya BioSciences) using a 20X 0.8 NA air objective (Olympus). Requisite solutions for this instrument include ACS-grade DMSO (Fisher Chemical), nuclease-free water, and 1X PCF buffer with buffer additive, the latter of which was prepared by adding 100 mL of 10X PCF buffer and 100 mL of buffer additive to 800 mL of nuclease-free water. Further, low- and high-concentration DMSO solutions for the PhenoCycler Fusion 2.0 were prepared: low-concentration DMSO consisted of 10% DMSO and 10% additive buffer in nuclease-free water, whereas high-concentration DMSO consisted of 10% additive buffer and 10% nuclease-free water in DMSO.

Some antibody clones are listed below as AKYP clones rather than commercially available ones. The original clones as delivered were listed as available, but all new products are sold as manufacturer-specific. Second, some of the products are no longer available or now feature different barcode/reporter numbers. This list is representative of all products used in this study.

| **PCF Antibody** | **Clone** | **Barcode/Reporter** | **Wavelength** | **Catalog Number** | **Exposure Time (ms)** |
| --- | --- | --- | --- | --- | --- |
| CD20 | L26 | BX/RX020 | AF750 | 4450018 | 225 |
| CD8 | C8/144B | BX/RX026 | Atto550 | 4250012 | 150 |
| CD4 | EPR6855 | BX/RX003 | AF647 | 4550112 | 180 |
| GZMB | D6E9W | BX/RX041 | Atto550 | 4250055 | 108 |
| FOXP3 | 236A/E7 | BX/RX031 | AF647 | 4550071 | 225 |
| Ki67 | B56 | BX/RX047 | Atto550 | 4250019 | 180 |
| HLA-A | EP1395Y | BX/RX004 | AF750 | 4450046 | 180 |
| Galectin-3 | M3/38 | BX/RX035 | Atto550 | 4450034 | 90 |
| CD3e | EP449E | BX/RX045 | AF647 | 4550119 | 225 |
| CD45RO | UCHL1 | BX/RX017 | Atto550 | 4250023 | 300 |
| CD45 | D9M81 | BX/RX021 | AF647 | 4550121 | 50 |
| CD40 | AKYP0095 | BX/RX010 | AF647 | 4550064 | 150 |
| EPCAM | D9S3P | BX/RX091 | AF750 | 4450088 | 260 |
| PD-L1 | 73-10 | BX/RX043 | AF647 | 4550072 | 500 |
| CD14 | EPR3653 | BX/RX037 | Atto550 | 4450047 | 200 |
| PD-1 | D4W2J | BX/RX046 | AF647 | 4550038 | 400 |
| MPO | AKYP0113 | BX/RX098 | Atto550 | 4250083 | 150 |
| CD68 | KP1 | BX/RX015 | AF647 | 4550113 | 125 |
| IDO1 | V1NC3IDO | BX/RX027 | AF647 | 4550123 | 130 |
| CD31 | EP3095 | BX/RX001 | AF750 | 4150017 | 225 |
| KRT14 | Poly19053 | BX/RX002 | Atto550 | 4450031 | 25 |
| ICOS | D1K2T | BX/RX054 | AF647 | 4550117 | 125 |
| SMA | AKYP0081 | BX/RX013 | AF750 | 4450049 | 113 |
| PDPN | NC-08 | BX/RX023 | Atto550 | 4250094 | 125 |
| COL_IV | EPR20966 | BX/RX042 | AF647 | 4550122 | 170 |
| CD34 | AKYP0088 | BX/RX025 | Atto550 | 4250057 | 200 |
| HLA-DR | EPR3692 | BX/RX033 | AF647 | 4550095 | 90 |
| CD38 | E7Z8C | BX/RX089 | Atto550 | 4250080 | 100 |
| Bcl2 | EPR17509 | BX/RX085 | AF647 | 4250098 | 100 |
| VIM | O91D3 | BX/RX022 | AF750 | 4450050 | 70 |
| IFNG | AKYP0074 | BX/RX020 | Atto550 | 4250062 | 75 |
| PanCK | AE-1/AE-3 | BX/RX019 | AF750 | 4450020 | 170 |
| CD56 | CAL53 | BX/RX028 | Atto550 | 4250087 | 400 |
| CD11c | 118/A5 | BX/RX024 | AF647 | 4550114 | 60 |

<u>RNAscope HiPlex V2 on FFPE tissues</u>. All reagents were purchased from ACD and used as received unless otherwise noted according to our previous protocol^8^. In brief, slides were antigen retrieved as described above prior to undergoing the RNAscope protocol. After using an ImmEdge pen (Vector Laboratories, #H-400) to draw a hydrophobic barrier around the tissues, slides went through a hybridization process consisting of Protease III reagent, *16S* rRNA (#464461-T4) at 1:100 concentration, three amplification steps, and fluorophore hybridization. ACD-provided DAPI was added to each tissue for 30 s; then, the DAPI was tapped off the slides, and the slides were mounted using Prolong Gold Antifade (Invitrogen #P36930, Lot #2836845). Sample imaging was performed using a Leica DMi8 with THUNDER Imager (Leica Microsystems) using a 40× 1.3 NA oil objective.

Laser capture microdissection (LCM) of health, periodontitis and peri-implantitis biopsies.

Antigen Samples were sectioned onto PEN membrane slides (Thermo Fisher #LCM0522) and stored in a desiccator until use in experiments. Antigen retrieval was performed by immersing slides in xylene and decreasing concentrations of EtOH (100%, 95%, and 70%), followed by RNAse-free water. Then, using a water bath heated to 95 °C, samples were antigen retrieved for 20 min. Slides were then rinsed 3x in 1x RNAse-free PBS. Following this, the backs of the slides were wiped dry; then, the slides were stored in a desiccator until use. *As an important note, all antigen retrieval was performed the day of LCM*. Using an Arcturus Cellect LCM instrument (previously, Thermo Fisher Scientific; currently, Laxco, Inc.), we captured 1) gingival basal and suprabasal epithelial and 2) sub-epithelial stromal sections using a laser spot size of 7.5 µm and a laser power of 70 mW with a pulse duration of 5 ms. Sections acquired were no larger than 500 µm^2^. After the regions had been acquired, CapSure Macro LCM Caps (Thermo Fisher #LCM0211) were used to remove the laser-dissected region from the tissue. *As modifications to the manufacturer’s instructions*, we implemented two changes during the capture. First, to account for humidity effects on laser efficiency, in between sample sectioning, we returned the slides to the desiccator. Second, after the instrument presented the LCM caps, we removed the film containing the section using tweezers and stored the entire film in 1X PBS in a –80 °C freezer until whole-genome sequencing of the FFPE sections was completed.

<u>DNA isolation from LCM FFPE sections.</u> Samples were transferred to a 2 mL tube containing 200 mg of 106/500 µm glass beads (Millipore-Sigma, St. Louis, MO) and 0.6 mL of Qiagen ATL buffer (Hilden, Germany). The suspension was agitated for 10 min on a digital vortex mixer at 3000 rpm and supplemented with 60 mg/mL lysozyme (Thermo Scientific, Rockford, IL). The suspension was incubated at 37 °C for 1 h and supplemented with 600 IU of Qiagen proteinase K, followed by an overnight 55 °C incubation. After incubation, the solution was centrifuged for 3 min, and 0.5 mL of supernatant and 0.5 mL of Qiagen AL buffer were incubated at 70 °C for 10 min. Then, the supernatant was aspirated and transferred to a new tube containing 0.5 mL of ethanol. DNA was purified using a standard on-column purification method with Qiagen buffers AW1 and AW2 as washing agents and eluted in DNase-free water^50,51^.

*<u>16S</u>* <u>rRNA amplicon sequencing.</u> 12.5 ng of total DNA were amplified using universal primers targeting the V4 region of the bacterial *16S* rRNA gene. Primer sequences contained overhang adapters appended to the 5’ end of each primer for compatibility with the Illumina sequencing platform. The primers used were F515/R806. Master mixes contained 12.5 ng of total DNA, 0.5 µM of each primer, and 2x KAPA HiFi HotStart ReadyMix (KAPA Biosystems, Wilmington, MA). The thermal profile for the amplification of each sample had an initial denaturing step at 95 °C for 3 min, followed by cycling of denaturing at 95 °C for 30 s, annealing at 55 °C for 30 s, and a 30-s extension at 72 °C (25 cycles), a 5-min extension at 72 °C and a final hold at 4 °C. Each *16S* amplicon was purified using the AMPure XP reagent (Beckman Coulter, Indianapolis, IN). Next, each sample was amplified using a limited cycle PCR program, adding Illumina sequencing adapters and dual-index barcodes (index 1(i7) and index 2(i5)) (Illumina, San Diego, CA) to the amplicon target. The thermal profile for the amplification of each sample had an initial denaturing step at 95 °C for 3 min, followed by cycling of denaturing at 95 °C for 30 s, annealing at 55 °C for 30 s, and a 30-s extension at 72 °C (25 cycles), a 5-min extension at 72 °C and a final hold at 4 °C. The final libraries were again purified using the AMPure XP reagent (Beckman Coulter), quantified, and normalized before pooling. The DNA library pool was then denatured with sodium hydroxide, diluted with hybridization buffer, and heat-denatured before loading on the MiSeq reagent cartridge (Illumina) and the MiSeq PE250 instrument (Illumina). Automated cluster generation and paired-end sequencing with dual reads were performed according to the manufacturer’s instructions^52,53^.

<u>Whole-genome sequencing (WGS).</u> 5 ng of genomic DNA were processed using the Nextera XT DNA Sample Preparation Kit (Illumina). Target DNA was simultaneously fragmented and tagged using the Nextera Enzyme Mix containing transposome that fragments the input DNA and adds the bridge PCR (bPCR)-compatible adaptors required for binding and clustering in the flow cell. Fragmented and tagged DNA was amplified and purified as described in the previous section. The DNA library pool was loaded on the Illumina platform reagent cartridge (Illumina) and the NextSeq 2000P4 platform (Illumina).

### Bioinformatics analyses of laser capture microdissection (LCM) data

<u>Output analysis of WGS data.</u> Sequencing output from the Illumina NextSeq 2000P4 platform were converted to fastq format and demultiplexed using Illumina BCL Convert 4.2.4. Quality control of the demultiplexed sequencing reads was verified by FastQC. Adapters were trimmed using Trim Galore. The resulting paired-end reads were submitted to Kraken2 for taxonomic classification^54^. An estimate of taxonomic composition including host was produced from these results using Bracken 2.5^55^. All reads classified as host were eliminated and paired-end reads were joined with VSEARCH 2.7.0^56^. Estimates of taxonomic composition, gene family, path abundance, and path coverage were produced from the remaining reads using HUMAnN3^57^. Diversity analysis was performed using QIIME2^58^.

<u>Output analysis of *16S* rRNA amplicon sequencing.</u> Data were converted to .fastq format and demultiplexed using the Ilumina Bcl2Fastq 2.20.0. The resulting paired-end reads were processed using QIIME 2 2024.5. Index and linker primer sequences were trimmed using the QIIME 2 invocation of Cutadapt. The resulting paired-end reads were processed with DADA2 through QIIME 2 including merging paired ends, quality filtering, error correction, and chimera detection. Amplicon sequencing units from DADA2 were assigned taxonomic identifiers based on the Silva 138 database using the QIIME 2 q2-feature classifier. Alpha diversity indexes Faith PD whole tree, Evenness (Shannon), and observed species were estimated using QIIME 2 at a rarefaction depth of 5,000 sequences. Beta diversity estimates were calculated within QIIME 2 using weighted and unweighted Unifrac distances and Bray-Curtis dissimilarity between samples at a subsampling depth of 5,000. The significance of differential abundance was estimated using ANCOM as implemented in QIIME 2^59–61^.

<u>DNA isolation process validation.</u> For validation of the DNA isolation process, a known bacterial community, ZymoBIOMICS Gut Microbiome Standard (Cat #D6331), and blanks composed of only DNA isolation reagents were included in the DNA extraction process and again in the library preparation. In addition to the isolation controls, the library preparation included library blanks composed of library preparation reagents alone.

<u>Diversity and differential abundance analyses of WGS data.</u> Alpha diversity was assessed using four indices: Shannon, Simpson, richness, and evenness. Beta diversity was evaluated using the first two principal coordinates from a Principal Coordinates Analysis (PCoA) based on Bray– Curtis dissimilarity. For differential abundance analysis, microbial count data were normalized using counts per million (CPM), and a log-normal model was applied to identify species associated with condition, adjusting for anatomical niche (epithelial and stromal compartments).

### Computational analyses of imaging data

<u>Cell segmentation for mIF and RNAscope data using Cellpose:</u> We used an extension for Cellpose within QuPath to segment the health, periodontitis, peri-implantitis, Grade C periodontitis, and matched healthy sample data, acquired from PCF or RNAscope, with the results exported as a .txt file. These segmentation results were used for further analyses.

<u>TACIT (Threshold-based Assignment of Cell Types from Multiplexed Imaging Data) for cell type</u> <u>annotation of imaging data.</u> After cell segmentation, TACIT was applied to resolve cell identities from all mIF-derived data^20^. TACIT enabled precise cell detection, separating predefined cell type signatures from background noise using a cell type relevance deconvolution algorithm.

<u>Constellation for tissue cellular neighborhood assignment in whole slide imaging.</u> Following cell assignment of mIF data using TACIT, Constellation was applied to extract tissue cellular neighborhoods (TCNs) from the whole-slide imaging data^7^. Using a consensus-based assignment approach, the TACIT data was partitioned into biologically relevant TCNs based on likelihood of cell occurrence and spatial prediction.

## DATA AVAILABILITY

All data, including links to the original raw data (GSE152042, GSE161267, GSE164241, GSE266897) can be found at: https://cellxgene.cziscience.com/collections/71f4bccf-53d4-4c12-9e80-e73bfb89e398. Additional raw data from the peri-implantitis single-cell RNA sequencing samples is under upload to GEO.

## CODE AVAILABILITY

Analysis notebooks are available at https://github.com/loci-lab/CD38-vasculopathy.

## ACKNOWLEDGEMENTS

We would like to acknowledge the NIH/NIDCR Dental Clinic, specifically Rachel Adam, Danielle Elangue, and Janice Lee, for their continued support to provide human tissues. Services in support of this work were provided by the VCU Massey Comprehensive Cancer Center Bioinformatics Shared Resource, supported in part with funding from the NIH-NCI Cancer Center Support Grant (#P30CA016059). The UNC Microbiome Core is funded in part by the Center for Gastrointestinal Biology and Disease (CGIBD P30 DK034987) and the UNC Nutrition Obesity Research Center (NORC P30 DK056350). This work was supported by the Chan Zuckerberg Initiative (Next-Gen Researcher Pilot Award) to QTE, the Fundação de Amparo à Pesquisa do Estado de São Paulo (FAPESP; #2021/11082-4) to CSS, Nihon University (2022 Overseas Researcher Grant) to AH, the National Institute of Dental and Craniofacial Research (#1R03DE034507-01) to DW, the Penn School of Dental Medicine Oral Maxillofacial Surgery Department (Schoenleber Grant) to KIK, and generous start-up funds from the ADA Science & Research Institute (Volpe Research Scholar Award) and the Philips Institute for Oral Health Research, as well as the American Academy of Implant Dentistry Foundation (Large Research Grant) to KMB. This publication is part of the Human Cell Atlas: https://www.humancellatlas.org/publications

## AUTHOR CONTRIBUTIONS

For this study, KMB and QTE conceptualized the project. QTE, KLAH, JX and AH developed methods for project analysis. Janice Lee (National Institute of Dental, Oral & Craniofacial Research), RC, and KIK supported the recruitment of patients and collected data. QTE, CSS, ZC, KIK, RC & KMB collected samples for analysis. QTE, KLAH, JX, BFM, AH, ZAM, AAR, NP, AMA-P & KMB performed experimental and/or bioinformatic analysis. QTE and KMB wrote the original draft; all authors critically reviewed and edited the final manuscript.

---

**Figure 1S.**
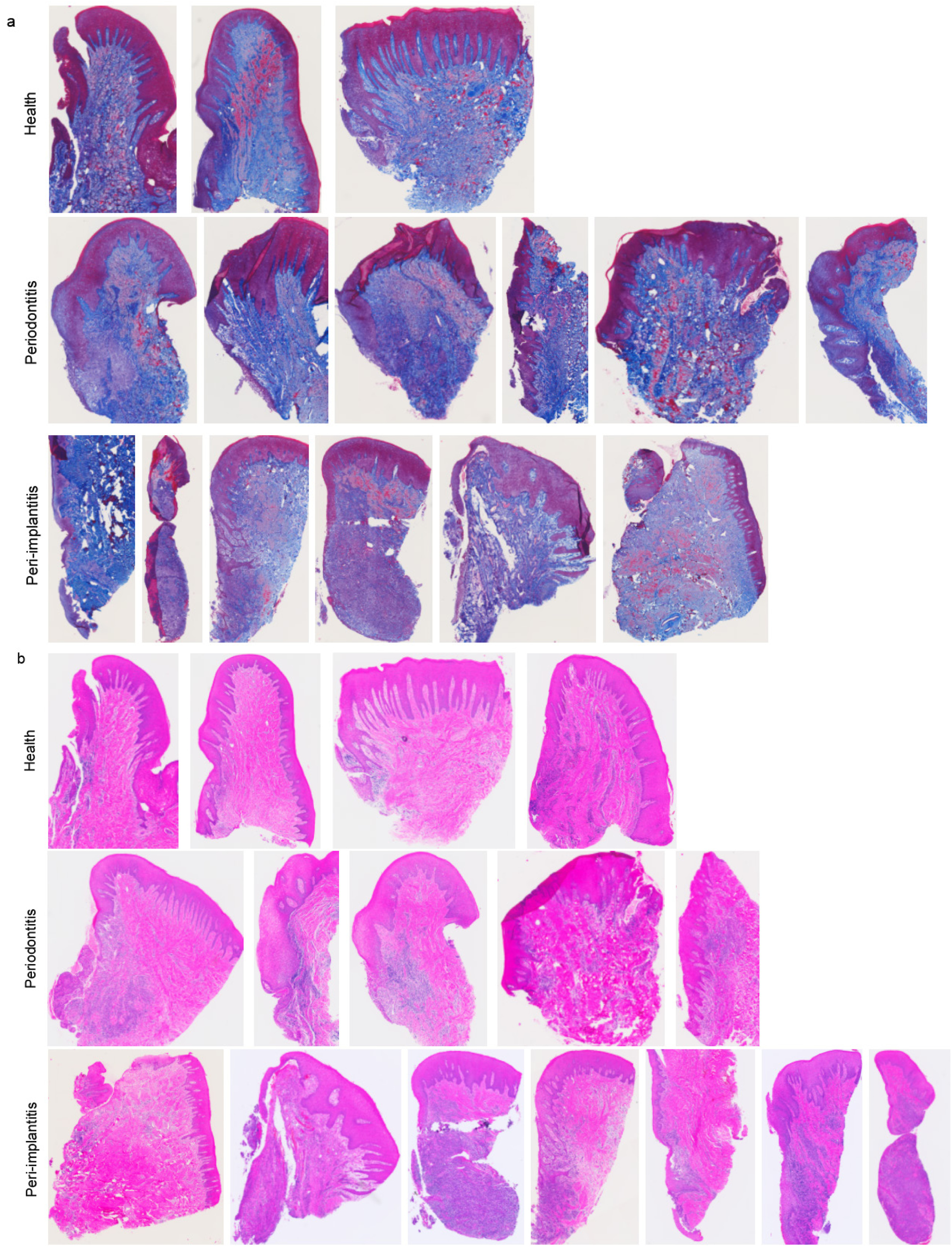

**Figure 2S.**
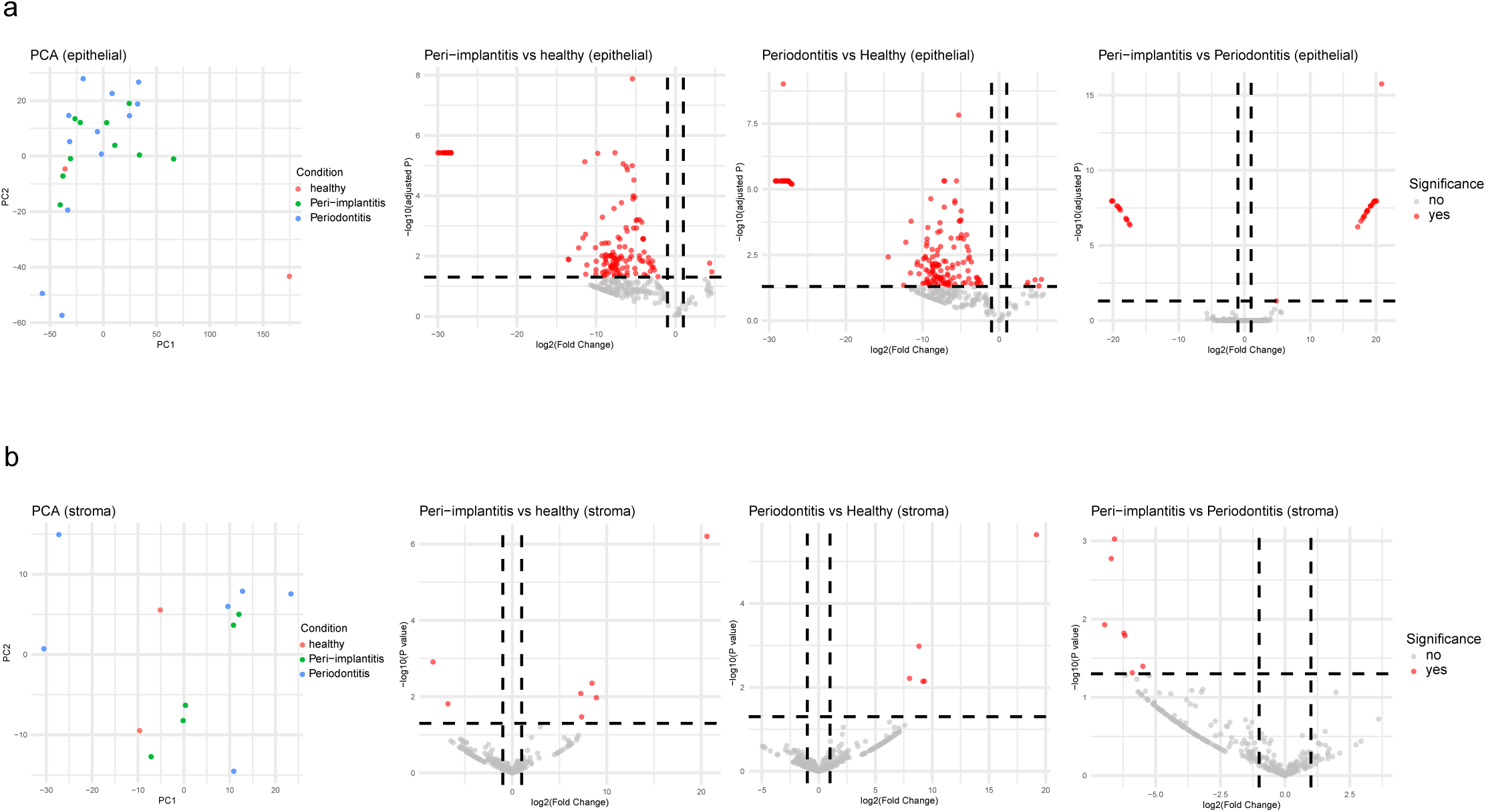

**Figure 3S.**
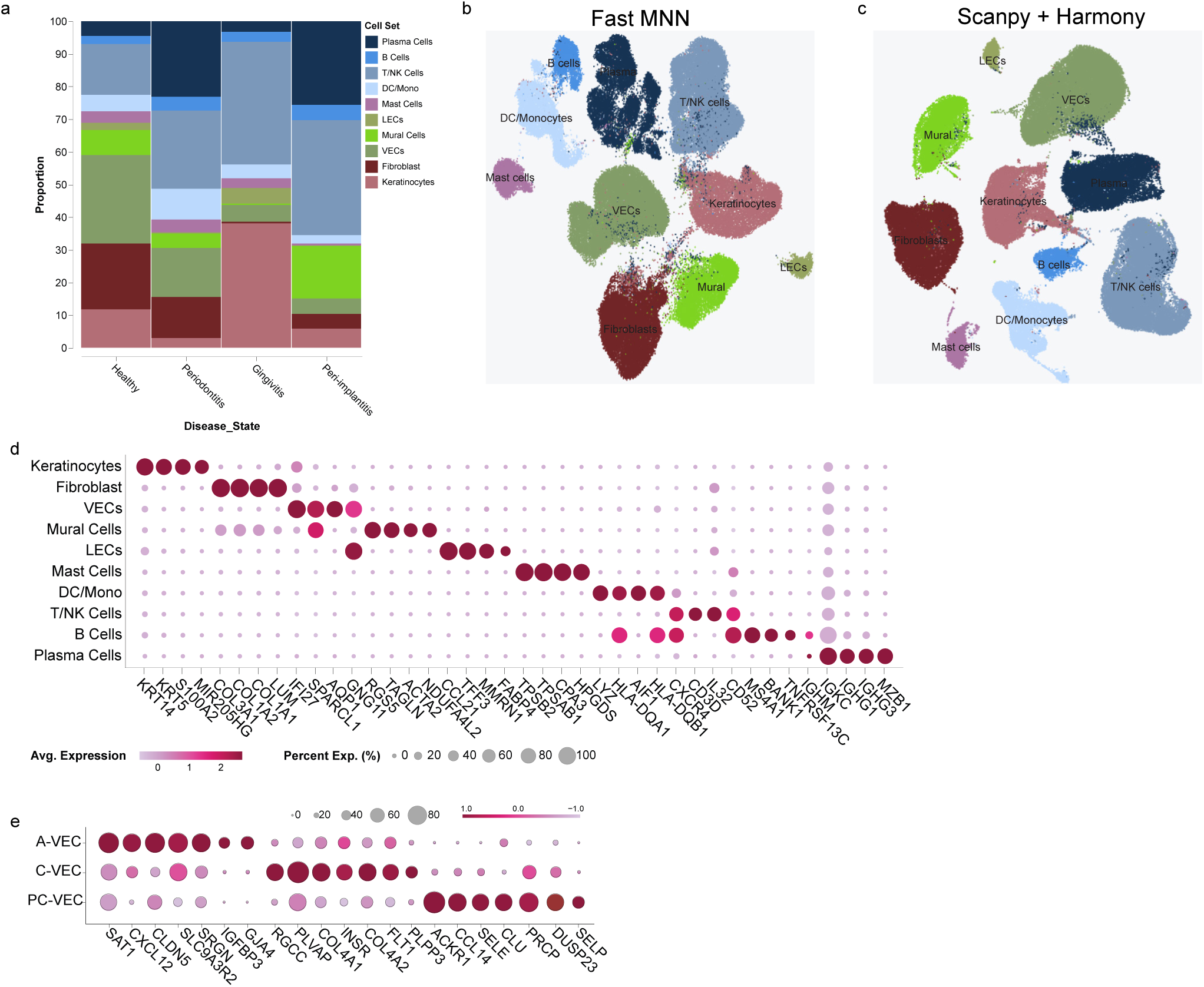

**Figure 4S.**
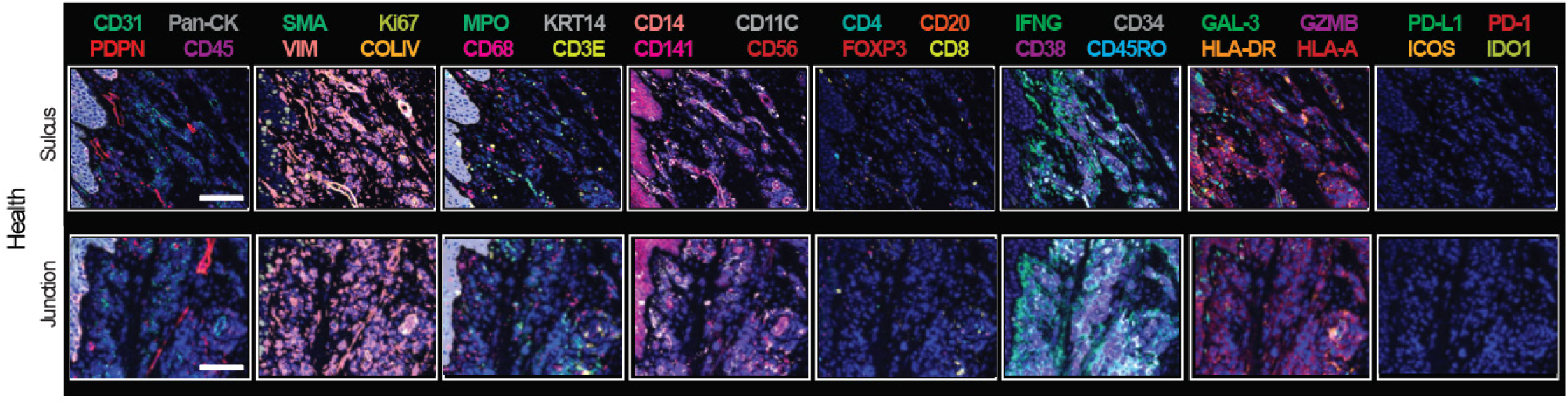

**Figure 5S.**
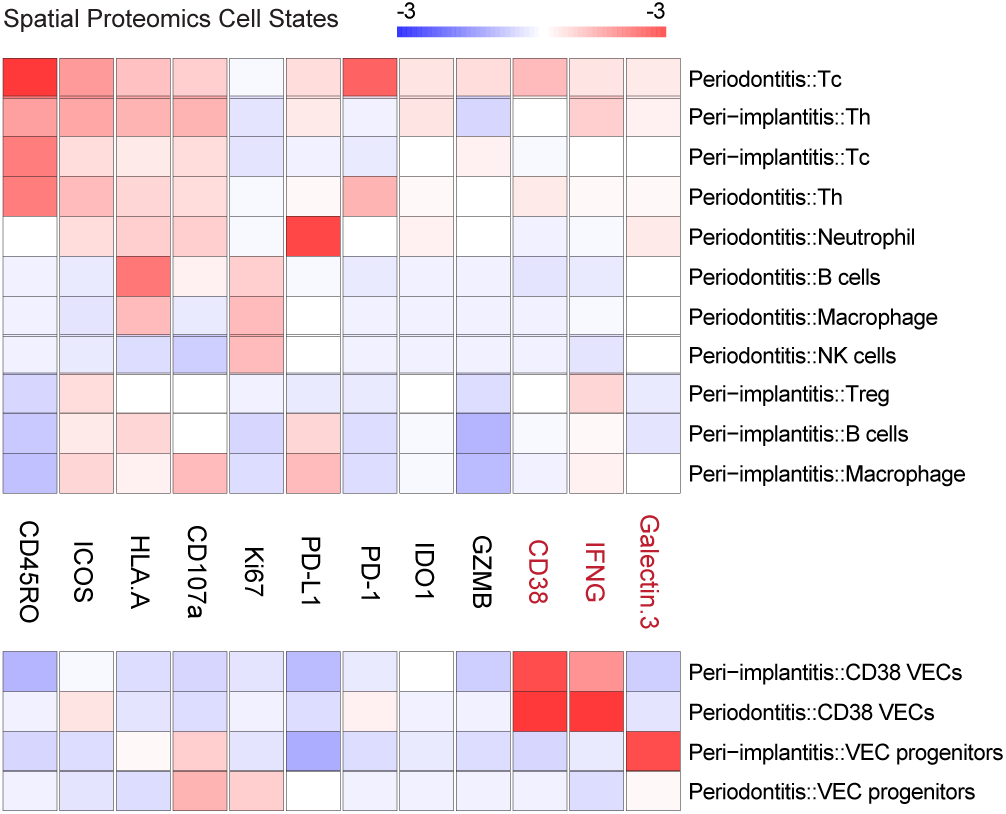

