## Supplemental Data 1 for "CD38^+^ Endothelial Remodeling Defines Spatially Diverse Vasculopathy Programs in Rapidly Advancing Oral Inflammation"

**SUPPLEMENTARY DATA DESCRIPTIONS**

**Supplementary Figures**

Figure 1S. a) Masson’s and b) H&E images of health, periodontitis, and peri-implantitis biopsies.

Figure 2S. a) Epithelial- and b) stromal-specific alpha- and beta-diversity analyses.

Figure 3S. a) Proportions of cell types per condition. b,c) Additional clustering types. Differentially expressed gene plots for d) all and e) vascular endothelial cell subpopulations.

Figure 4S. Healthy mIF data acquired from PhenoCycler Fusion.

Figure 5S. Spatial proteomics cell states in periodontitis and peri-implantitis.

**Supplementary Tables**

Supplementary Table 1. Metadata for health, periodontitis, peri-implantitis, Grade C periodontitis, and matched healthy samples.

Supplementary Table 2. Differentially expressed gene lists for Tier 1 cell sets and Tier 2 vascular endothelial cell subpopulations.
